# Ultrasound pulse repetition frequency preferentially activates different neuron populations independent of cell type

**DOI:** 10.1101/2024.03.25.586645

**Authors:** Jack Sherman, Emma Bortz, Erynne San Antonio, Hua-an Tseng, Laura Raiff, Xue Han

**Affiliations:** Department of Biomedical Engineering, Boston University; Department of Pharmacology and Experimental Therapeutics, Boston University

**Author notes:** equal contribution (Jack Sherman, Emma Bortz).

**Keywords:** Transcranial ultrasound, neuromodulation, GCaMP7f, mechanosensitive ion channels, parvalbumin interneurons, excitatory neurons

## Abstract

Transcranial ultrasound activates mechanosensitive cellular signaling and modulates neural dynamics. Given that intrinsic neuronal activity is limited to a couple hundred hertz and often exhibits frequency preference, we examined whether pulsing ultrasound at physiologic pulse repetition frequencies (PRFs) could selectively influence neuronal activity in the mammalian brain. We performed calcium imaging of individual motor cortex neurons, while delivering 0.35 MHz ultrasound at PRFs of 10, 40, and 140 Hz in awake mice. We found that most neurons were preferentially activated by only one of the three PRFs, highlighting unique cellular effects of physiologic PRFs. Further, ultrasound evoked responses were similar between excitatory neurons and parvalbumin positive interneurons regardless of PRFs, indicating that individual cell sensitivity dominates ultrasound-evoked effects, consistent with the heterogeneous mechanosensitive channel expression we found across single neurons in mice and humans. These results highlight the feasibility of tuning ultrasound neuromodulation effects through varying PRFs.

## Introduction

Transcranial low-intensity ultrasound (US) neuromodulation represents a promising therapeutic strategy for neurological and psychiatric disorders^1,2^. Noninvasive US neuromodulation typically uses lower fundamental frequencies of hundreds of kilohertz to a couple megahertz due to superior skull penetration ^3–8^. In addition to fundamental frequency and acoustic pressure, various permutations of US pulse repetition frequencies (PRFs), durations, and intervals create an essentially infinite parameter space.

As a mechanical energy wave, US interacts with cellular membranes through various acoustic phenomena which could induce phospholipid reconfiguration, alterations in membrane fluidity and permeability^9–13^, or changes in membrane curvature. Though the exact physical interactions between US and cellular membranes remain unknown, mechanosensitive channels are generally thought to be critical in mediating cellular signaling changes during US neuromodulation ^14–19^. Many mechanosensitive channels are widely expressed in the brain ^20^, such as the two-pore potassium K2P family ^20–22^, hyperosmolality-gated calcium-permeable TMEM63 family ^20,23^, mechanosensitive cation channel Piezo family ^20,24,25^, and the transient receptor potential TRP families ^19,20,26–30^. Among the large number of mechanosensitive channels, several members of the K2P, TRP and Piezo1 families have been implicated in ultrasound neuromodulation ^19,24–26,31,32^ (**Supplemental Table 1**). For example, genetic knockout of Piezo1^25^ or knockdown of Trpm2 ^26^ reduced US-mediated neural and behavioral responses in mice. Similarly, genetic knockdown of TRPC1, TRPP1, and TRPP2 attenuated cellular calcium responses in cultured neurons *in vitro*^19^.

While mechanosensitive channels are broadly distributed in brain tissue, it is unclear how mechanosensitive channel expression differs between individual cells. In this study, we first analyzed Allen Brain Institute’s single cell sequencing database and confirmed extensive variation of mechanosensitive channel expression among individual neurons in both mice and humans. Interestingly, some channels show stronger expression in excitatory neurons than inhibitory neurons, though the differences between cell types are much less pronounced than between individual neurons. Such systemic variation among cell types could result in distinct mechano-sensitivity and thus differential responses between cell populations to US.

The effect of US not only depends on the direct activation of individual neurons but also on subsequent network responses in intact brain circuits. At a fundamental frequency of around 0.5 MHz, typically used for transcranial neuromodulation, the theoretical focal spots are several millimeters in diameter, thus recruiting a large tissue volume. This lack of focus at lower US fundamental frequencies is particularly prominent when studying US effects in mouse models with a brain size of about 13×11×8 mm^3^, where about a quarter or more of the brain is more or less directly sonicated. Indeed, experimental studies confirmed that US could produce both excitatory and inhibitory neuronal changes in humans ^3–7,33^ and animal models^25,26,34–37^. For example, 0.5 MHz US delivered at 1 kHz PRF suppressed somatosensory evoked potentials in humans^33^, whereas 0.25 MHz US at 500 Hz PRF excited the somatosensory cortex ^5^. Our previous study also confirmed that ultrasound evoked more prominent cellular calcium responses in the motor cortex than the hippocampus in awake mice^35^. Despite the difficulty of dissociating direct US-evoked cellular effects from indirect network responses, US at 900 Hz PRF was recently shown to activate parvalbumin expressing interneurons at a population level, while suppressing the excitatory neuron population in the mouse hippocampus ^34^. Similarly, US pulsed at 30 Hz, 300 Hz, and 1.5 kHz PRFs produced slightly different responses in excitatory versus inhibitory neuron populations^37^.

Some mechanosensitive channels, such as TRPs and Piezo1, are permeable to cations including calcium, and thus their activation depolarizes the plasma membrane and increases intracellular calcium ^19,24–26^. Both membrane depolarization and intracellular calcium rise could engage downstream signaling that leads to increased spiking probability. Since US neuromodulation generally exhibits weak effects, the intracellular pathways downstream of mechanosensitive channel activation, in particular activation of voltage-gated sodium and potassium channels critical for action potential generation, likely provide additional amplification mechanisms to translate acoustic driven responses into spiking outputs ^14–17,19^. This is supported by the observation that US activation of mechanoreceptors TRPP and TRPC led to intracellular calcium increases, which subsequently recruit calcium-activated TRPM4 channels and voltage-gated T-type calcium channels to increase spiking probability in cultured neurons ^19^.

The kinetics of neuronal ion channels generally limit the speed of spiking and subthreshold membrane voltage fluctuations to a couple hundred hertz. Many neurons have intrinsic biophysical properties that render them more sensitive to stimulation at certain physiologic frequencies of a few hertz to tens of hertz. We recently demonstrated that when electrical stimulation frequency reaches 140 Hz, membrane voltage was poorly paced by individual electrical pulses ^38^. Thus far, most studies have delivered US at PRFs on the order of kilohertz. However, it is highly plausible that pulsing US at physiological frequencies could optimize the coupling of mechanosensitive channel activation to subsequent signaling pathways, which together could improve US modulation efficiency.

To examine whether US at different physiological PRFs selectively influence neuronal response in the awake mammalian brain, we performed large-scale single cell calcium imaging from parvalbumin-positive interneurons and parvalbumin-negative predominantly excitatory neurons in awake head-fixed mice while pulsing 0.35 MHz US at 10 Hz, 40 Hz, or 140 Hz. We chose these frequencies based on previous observations that they likely engage biophysical signaling differently in motor cortex neurons. 10 Hz is within the beta frequency band, an intrinsically preferred frequency of the cortical-basal ganglia motor circuits, and readily supported by the motor cortex network ^39^. 40 Hz is within the gamma frequencies (∼30-100 Hz) that have been broadly linked to PV interneuron oscillations ^40,41^. Transcranial ultrasound pulsed at 40 Hz augmented gamma oscillations ^42^ and reduced amyloid beta in mouse models of Alzheimer’s disease ^42,43^. Further, PV cells were implicated in the 40 Hz visual stimulation induced gamma oscillations and amyloid beta reduction ^44^. 140 Hz is widely used in deep brain stimulation, which leads to better therapeutic outcomes than lower frequency stimulations. Our recent studies demonstrated that 140 Hz electrical stimulation robustly depolarized membrane voltage, scrambled spike timing and led to informational lesion ^38^

Consistent with our analysis of the Allen Brain Institute’s single cell sequencing results showing prevalent and heterogeneous expression of many mechanosensitive channels across individual neurons, we detected diverse US-mediated cellular calcium responses across neurons, with many reliably activated. US-evoked calcium events in individual neurons exhibit the same characteristics as those naturally occurring during increased spiking probability, suggesting that US increases neuronal activity. By calculating pair-wise correlations between simultaneously recorded neurons, we confirmed that US transiently increased network synchrony without producing prolonged over-synchronization that may be detrimental to neurological functioning. Further, by directly comparing the evoked responses to different PRFs in the same neuron, we revealed that most neurons were preferentially activated by a single PRF. Because of the frequency preference of individual cells, different populations of neurons were selectively recruited by specific PRFs. Finally, the evoked population responses across all PRFs tested were similar between the two neuron types, highlighting the importance of considering individual neuron heterogeneity regardless of conventionally defined cell type.

## Results

### Mechanosensitive channel expression is prominent and heterogeneous across individual neurons in the cortex and hippocampus of mice and humans

Mechanosensitive channels are critical in translating mechanical forces to cellular responses, and many members of the K2P, TRP, TMEM and Piezo families of mechanosensitive channels are widely expressed in the brain (Supplemental **Table 1**). However, there has been no systematic analysis of the expression patterns of these channels across brain regions or cell types. We first analyzed the Allen Mouse Brain Atlas in-situ hybridization gene mapping data to examine their brain-region dependent expression ^45^ (Supplemental **Table 2,** https://mouse.brain-map.org). We found that many mechanosensitive channels are widely expressed throughout the brain, and the expression levels for some channels are more variable than others. TRPC1, TRPM2&7, TMEM63B, and Piezo1 exhibit high expression levels throughout the brain, with some variations across brain regions. In contrast, Trpm3 expression is more restricted to the hippocampus, olfactory areas, and cerebellum. Such variation may underlie the reported difference in US mediated effects across brain regions ^35^.

To further evaluate single cell level expression of these mechanosensitive channels, we analyzed Allen Brain Institute’s single-cell RNA-sequencing transcriptome databases containing 76,533 cells throughout the human motor cortex and 1.1 million cells throughout the mouse cortex and hippocampus ^46–48^. We found that many channels are widely expressed across transcriptomically defined cell types in humans (**Fig. 1a**) and in mice (**Fig. 1b**). Interestingly, in humans, KCNK2 and TRPC6 appear to have stronger expression in inhibitory neurons, while KCNK10 are expressed predominantly in excitatory neurons (**Fig. 1a, Supplemental Table 3**). Similarly, in mice, some channels, including TRAAK, TRPM3, TREK2, and TRPC1, have significantly higher expression levels in excitatory neurons (Slc17a7 positive) than inhibitory neurons (GAD1 positive) in mice (**Fig. 1b**, **Supplemental Table 4**). Curiously, Piezo1, previously demonstrated to play a role in mediating US modulation ^24,25^, appeared negative in the transcriptome database, even though it is expressed across brain regions based on the in-situ hybridization gene mapping data (**Table 2**). This may be due to Piezo1 expression being concentrated in a subset of neurons, as the transcriptome database uses the trimmed mean resulting a gene to be negative if it is expressed in <25% of a given cluster. Thus, we further analyzed Piezo1 expression in the more recent whole brain ABC atlas (https://knowledge.brain-map.org/abcatlas) and confirmed its expression in a small fraction of both excitatory and inhibitory neurons.

**Figure 1:**
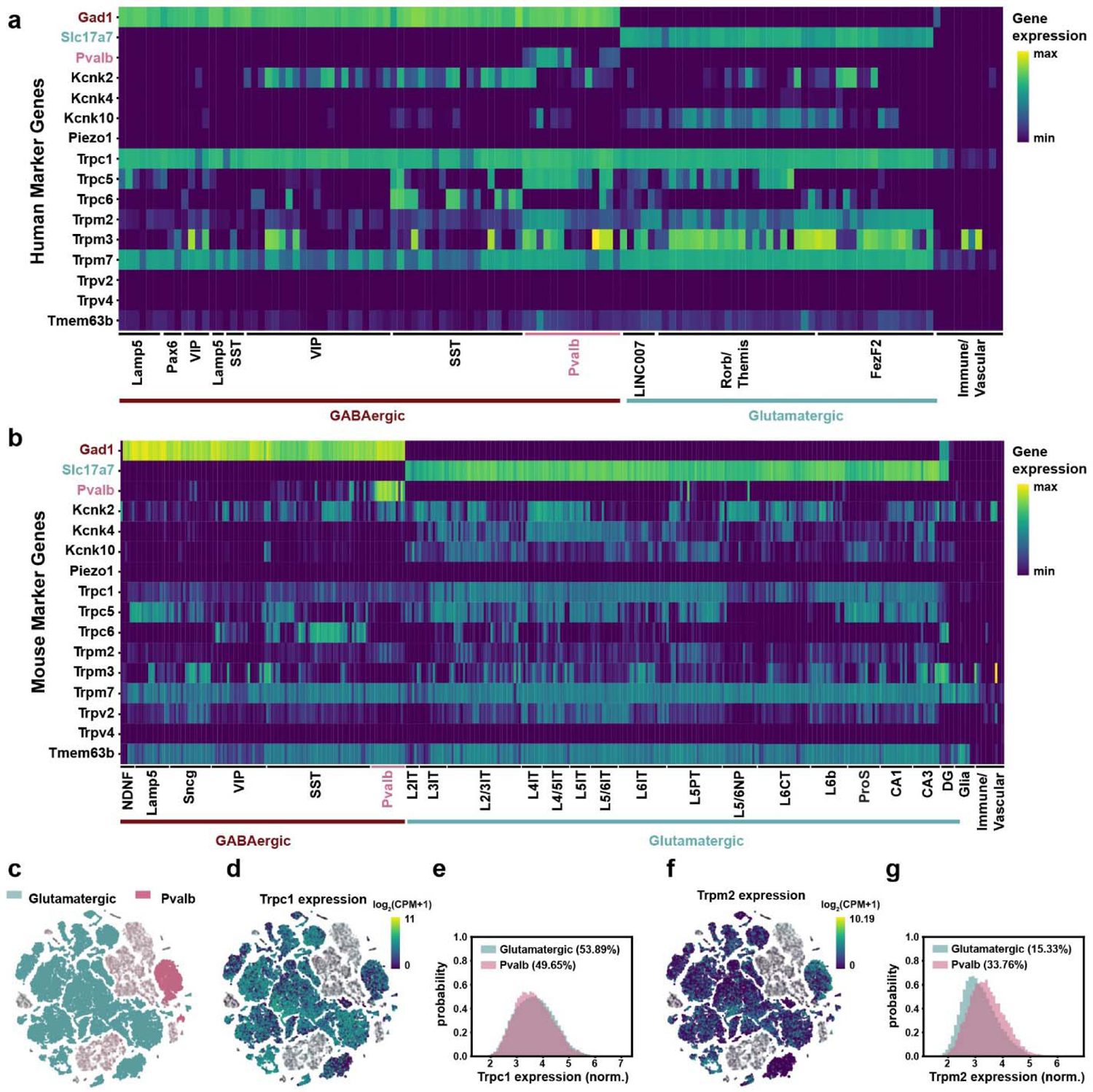
Widespread and heterogeneous expression of mechanosensitive channels in human and mice brains. (**a,b**) Heatmap of mechanosensitive ion channel gene expression in the human motor cortex (**a**) and the mouse motor cortex and hippocampus (**b**). Relative expression levels of example canonical marker genes and selected mechanosensitive channel genes are shown across common cell types within GABAergic and Glutamatergic neuron classes. Cell types are organized by hierarchical clustering. Gad1, inhibitory neuron marker gene; Slc17a7, excitatory neuron marker gene; and Pvalb, parvalbumin. Data are from Allen Institute for Brain Science, https://brain-map.org/atlases-and-data/rnaseq/human-m1-10x and https://brain-map.org/atlases-and-data/rnaseq/mouse-whole-cortex-and-hippocampus-10x. (**c, d, f)** Uniform Manifold Approximation and Projection (UMAP) representation of the mouse single cell sequencing results, colored by cell types (**c**, glutamatergic, teal; Pvalb, pink; Other subtypes: gray), overlapped with Trpc1 expression (**d**) or Trpm2 expression (**f**). Expression was scaled by log_2_(CPM+1), and the graphs were created with Cytosplore Viewer, https://viewer.cytosplore.org/. (**e,g**), Histograms of normalized expression of Trpc1 (**e**) and Trpm2 (**g**) in glutamatergic and PV neurons. Only cells with expression were included in probability density calculation, as shown by the percentage number in the legends. N = 30,461 PV cells and N = 976,358 glutamatergic cells.

Previous studies also reported differences in US-evoked responses between parvalbumin (PV) positive fast-spiking interneurons and excitatory neurons ^37^. Thus, we further compared gene expression patterns between PV positive interneurons and excitatory cells ^46–48^(Fig. 1c-g). We observed large variation in channel expression levels across individual neurons of both cell types, and the variation between individual cells was more prominent than between cell types. For example, expression distributions of Trpc1 and Trpm2, both implicated in ultrasound neuromodulation ^19,26^, varied significantly between PV interneurons and excitatory cell populations (**Fig. 1e,g**, Wilcoxon rank sum test, p < 0.01).

For Trpc1, a similar fraction (∼50%) of excitatory and PV cells had nonzero expression, and the expression level difference between the two populations was negligible (8 = −0.0363), indicating only a 3.6% probability that a glutamatergic neuron would have higher expression than a PV interneuron (**Fig. 1e**). For Trpm2, 34% of PV cells showed expression, in contrast to 15% of excitatory cells, and the expression level across the PV population was slightly higher than excitatory cells (**Fig. 1g**, 8 = 0.2315). Similar patterns are observed with the other channels examined, even though they are less explored in the context of US modulation (**Supplemental Fig. 1**). Despite excitatory neurons having higher expression of some mechanosensitive channels, the variation between individual neurons appears more prominent. Thus, the specific expression profiles of mechanosensitive channels in individual neurons are expected to have a greater impact on the outcome of ultrasound neuromodulation effect than canonical cell type.

### Calcium imaging analysis of ultrasound stimulation effect on spiking-related calcium activity in individual cortical neurons in awake mice

While the exact mechanisms supporting the transformation of US acoustic pressure to changes in cellular activity remain elusive, US-evoked effect is ultimately dictated by individual neuronal membranes’ biophysical properties and intracellular signaling ^14,15^. Most studies delivered US at high PRFs on the order of kilohertz, much higher than the maximum action potential rate neurons can support. Moreover, stimulation frequencies are also known to engage different neural activities. For example, in deep brain stimulation, pulse frequency is a critical consideration. We recently showed that 40 Hz electrical stimulation paced membrane potential and spike timing, but when electrical stimulation reached 140 Hz, spiking output failed to track stimulation pulses temporally ^38^. To gain a deeper understanding of how neurons cellular properties influence US-evoked activities, we analyzed the neuronal response to US delivered at biophysically relevant frequencies.

We performed single cell calcium imaging in awake mice using GCaMP7f ^49^, an improved genetically encoded calcium sensor with higher sensitivity for detecting spike-related calcium transients (Fig. 2a). We injected AAV9-syn-jGCaMP7f in the motor cortex of C57BL/6 mice, and then placed a glass imaging cranial window above the pia for optical access of GCaMP7f labeled neurons (Fig. 2b). During each experiment, awake mice were head fixed under a custom wide-field imaging microscope with an ultrasound transducer placed under the chin (Fig. 2a) delivering 350 kHz planar ultrasound.

**Figure 2:**
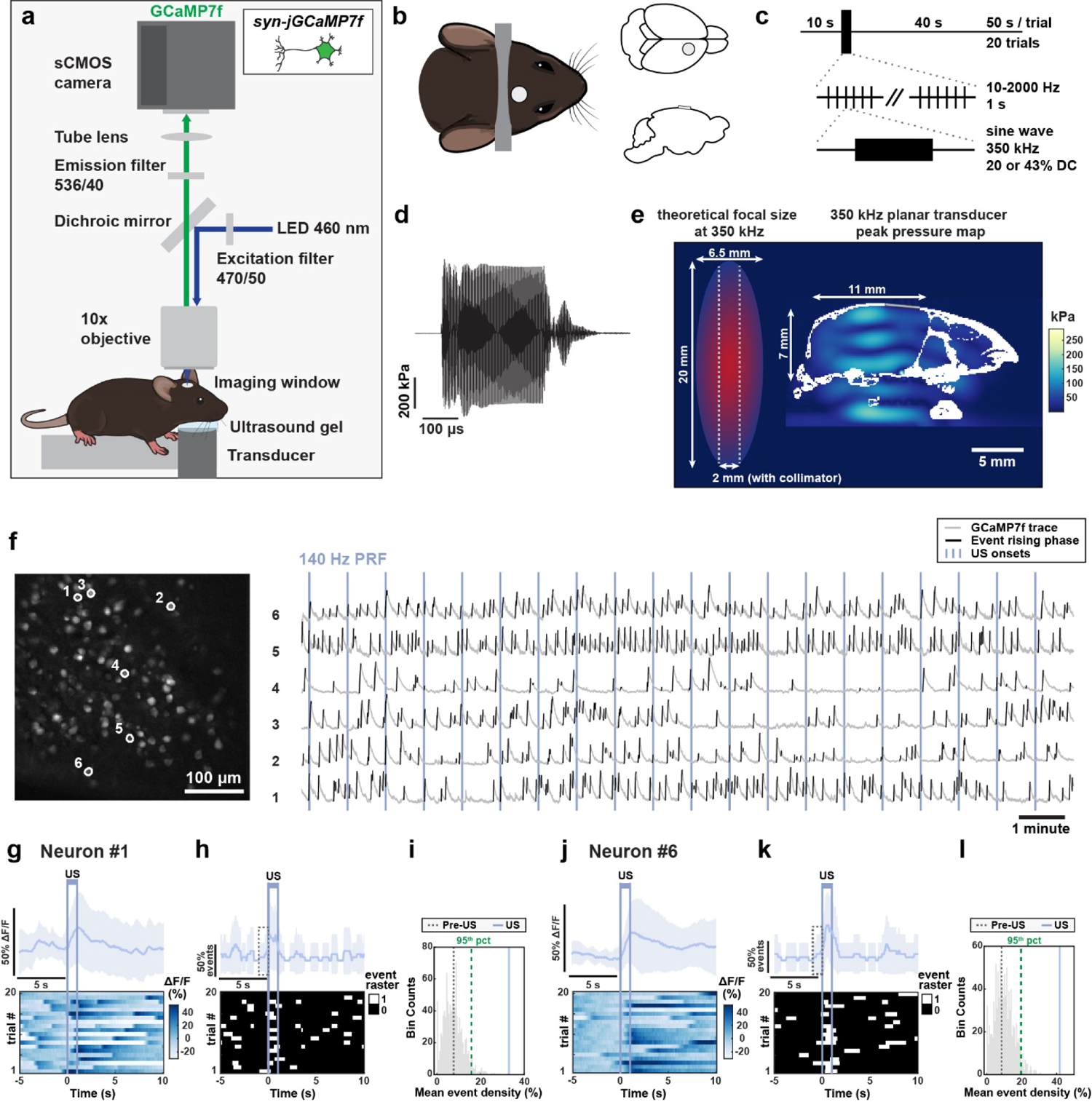
Cellular calcium imaging analysis of US-evoked calcium events associated with neuronal spiking in awake mice. (**a**). Experimental setup illustrating transcranial ultrasound stimulation on a head-fixed mouse during calcium imaging. The ultrasound transducer was placed beneath the mouse’s head and ultrasound gel acoustically coupled the transducer to the mouse. GCaMP7f expressing neurons were imaged with 460 nm LED excitation and 536/40 nm emission with an sCMOS camera. **(b)** Left: Imaging window and head bar placement, with the cranial window placed over the motor cortex. Right: Coronal and sagittal views of the cranial window. **(c)** Recording sessions consisted of 20 trials with 50 s each trial, containing a 10 s pre-US period followed by 1 s of 350 kHz US pulsed at 10, 40, 140, or 2000 Hz. **(d)** Example US waveform of the transducer recorded from an acoustic hydrophone in water. **(e)** Left: Theoretical −6dB dimensions of 350 kHz US focal spot and 2 mm diameter beam collimator, to scale with mouse head dimensions for acoustic pressure simulation. Right: Simulation of peak acoustic pressure in the mouse brain during 350 kHz, 522 kPa US. Skull geometry is highlighted in white and glass imaging window in gray. Scale bar is 5 mm. **(f)** Left: An example field of view’s max-min fluorescence projection image with example neurons circled in white. Scale bar = 100 µm. Right: GCaMP7f fluorescence traces from the six circled example neurons over 20 trials. GCaMP7f ΔF/F traces are in gray with detected calcium event rising phases highlighted in black. Vertical lines represent 140 Hz PRF US onset. (**g, j**) Top: Trial-averaged ΔF/F aligned to US onset for neuron #1 (**g**) and neuron #6 (**j**). Shading corresponds to SD over 20 trials. Bottom: Normalized GCaMP7f ΔF/F heatmap across trials. Vertical purple lines depict US onset and offset. (**h, k**), Top: Trial-averaged event density trace for neuron #1 (**h**) and neuron #6 (**k**). Shaded region represents SD over trials. Bottom: Binarized event rising phases over 20 trials. White represents timepoints with event rising phases and black represents timepoints with no event rising phases. Vertical lines represent US onset and offset. Dashed box depicts 1 s pre-US period. **(i, l)** Illustration of statistical shuffling test used to determine neuron #1 **(i)** and neuron #6 **(l)** being modulated. Shuffled event density during non-US “baseline” periods is shown in gray histogram, with the 95-percentile indicated by dashed green line. The observed event density during US is shown as a solid purple line, and the observed event density during the 1 s pre-US is shown as a dashed gray line.

Because of the small size of the mouse brain (13 mm × 14 mm × 8 mm), and a typical focused ultrasound radiation volume around ∼6.5 mm × 6.5 mm × 20 mm at 350 kHz, we chose to use planar ultrasound in this study (Fig. 2e). We placed the planar transducer under the chin to accommodate for the optics above the head for high-resolution single cell imaging. While thin ring transducers have been applied in mouse imaging studies^34,50^, their focusing ability is even more limited than standard focused ultrasound transducers. Similarly, collimators have been used to further limit the sonication area^51–53^ (Fig. 2e) but are too bulky for optical imaging compatibility. Because of the limited spatial resolution of sub-megahertz US, in this study we focused on deriving a principled understanding of how different physiologic PRFs engage distinct neurons, which could guide the selection of US pulsing protocols during clinical translation, rather than testing the effect of sonicating a particular neural circuit related to behavior or pathology.

While the mouth and nasal cavities shape the acoustic pressure profiles, US nonetheless propagates through the tissue, bone and skin to the motor cortex regions imaged. Our computational simulation estimated that the US delivered had a spatial peak pulse average intensity (I_SPPA_) of 0.2-3.16 W/cm^2^ at the imaging site. US was pulsed at physiologically relevant frequencies of 10 Hz, 40 Hz, or 140 Hz and the supra-physiological frequency of 2 kHz.

We filled the space between the ultrasound transducer and the mouse’s chin with ultrasound gel to allow for efficient acoustic wave propagation into the brain. Sham conditions used the same preparation without ultrasound gel. To assess whether US-evoked calcium responses are consistent with intrinsic physiological calcium events associated with spiking, we pulsed US for 1 second every 50 seconds, allowing us to capture sufficient intrinsic calcium events during the intertrial intervals. We recorded a total of 20 trials, resulting in a duration of 16.7-minutes per session (Fig. 2c).

To estimate the acoustic pressure intensity at the site of recording, we first recorded the spatial peak ultrasound pressure waveform in water with a needle hydrophone (Figure 2d). The acoustic pressure waveform had a peak amplitude of 522 kPa, corresponding to I_SPPA_ of 9.11 W/cm^2^ in water (Supplemental **Table 5-6**). We then used the recorded hydrophone pressure amplitude to computationally simulate the ultrasound pressure and intensity distributions in the mouse brain in k-Wave, as reported previously in Tseng et al (Fig. 2e, Supplemental **Table 6**). To approximate our experimental conditions, we modified the three-dimensional mouse skull from Chan et al, 2007 ^54^ to include a motor cortex craniotomy and a glass imaging window, and we added air pockets above and below the tongue to accommodate for below-the-chin transducer placement. Our computational models revealed that stimulating with 522 kPa ultrasound resulted in an *in situ* spatial peak pressure of 289.5 kPa (Supplemental **Table 6**). At the site of calcium imaging, the simulated average peak pressure ranged between 13-95 kPa and the pulse average intensity was 1.5 W/cm^2^. For computational model validation, we created a phantom mouse skull with the local bone structure above the motor cortex removed for hydrophone access. We then filled the phantom with ultrasound gel and measured pressure with a needle hydrophone at the approximate site of motor cortex calcium imaging. The measured pressure amplitude in the mouse skull during stimulation was ∼125 kPa, consistent with the upper bounds of our simulated estimate.

To examine US effects on individual neurons, we extracted neuronal GCaMP7f fluorescence traces over time (Fig. 2f) and identified calcium events as those with large amplitude increases in GCaMP7f fluorescence (see **Methods**). Across non-US periods, defined as the full recording duration excluding two seconds post-US, individual neurons had event rates of 3.08 ± 0.32 events/minute (mean ± SD, N = 16 imaging sessions in 9 mice), consistent with the general observation using GCaMP7f^55^. While the rising phase of a calcium event is related to spiking, the falling phase is attributed to both cytosolic calcium clearance and GCaMP7f’s calcium ion dissociation kinetics. Thus, to analyze US effect on spike-related calcium activity, we binarized GCaMP7f fluorescence traces with ones depicting calcium event rising phases and zeros elsewhere (Fig. 2g**-h**, Fig. 2j**-k**) and computed calcium event density, a measure of the fraction of time a neuron has heightened spiking probability.

During each recording session, we noticed that many neurons increased event density during US (Fig. 2g**-l**). To determine whether a neuron was activated by US, we compared the mean calcium event density during US (1 second interval) across all 20 trials (Fig. 2h, Fig. 2k) to the shuffled baseline event density distribution (Fig. 2i, Fig. 2l). To generate the shuffled baseline distribution for each neuron, we averaged the event rate over 20 randomly selected 1 second intervals during “baseline”, defined as the full recording period excluding 5 seconds after US onset, and then repeated this procedure 1000 times.

Neurons were deemed US-responsive if the event density during US was greater than the 95^th^ percentile of the corresponding shuffled baseline distribution. Because calcium event rising phase in the recorded neurons lasted for 1.11 ± 0.67 s (mean ± SD, N = 177,646 events), events starting in the second before US may lead to false identification. Thus, we excluded neurons that had event density rates higher than the 95^th^ percentile during the 1 second window prior to US.

### US delivered at physiologically relevant PRFs reliably increased the rates of calcium events associated with natural neuronal spiking

Of the 3,353 neurons imaged across all US parameters, 567 (16.9%) exhibited significantly increased calcium event density (Fig. 3a). US delivered at 10 Hz activated 162/861 neurons (18.82%) neurons, at 40 Hz activated 22/281 neurons (7.83%) neurons, and at 140 Hz activated 183/1019 neurons (17.96%) (Fig. 3a, Supplemental **Table 7**). When pulsed at the higher PRF of 2 kHz, US activated 200/1192 neurons (16.78%). As expected, in modulated neurons, the average event density rose immediately following US onset, while the event density remained unchanged in non-modulated neurons (Fig. 3b). The average event fluorescence waveform during US closely matched the average event density waveform during non-US periods, indicating that US induces naturalistic calcium events related to neuronal spiking (Fig. 3c).

**Figure 3:**
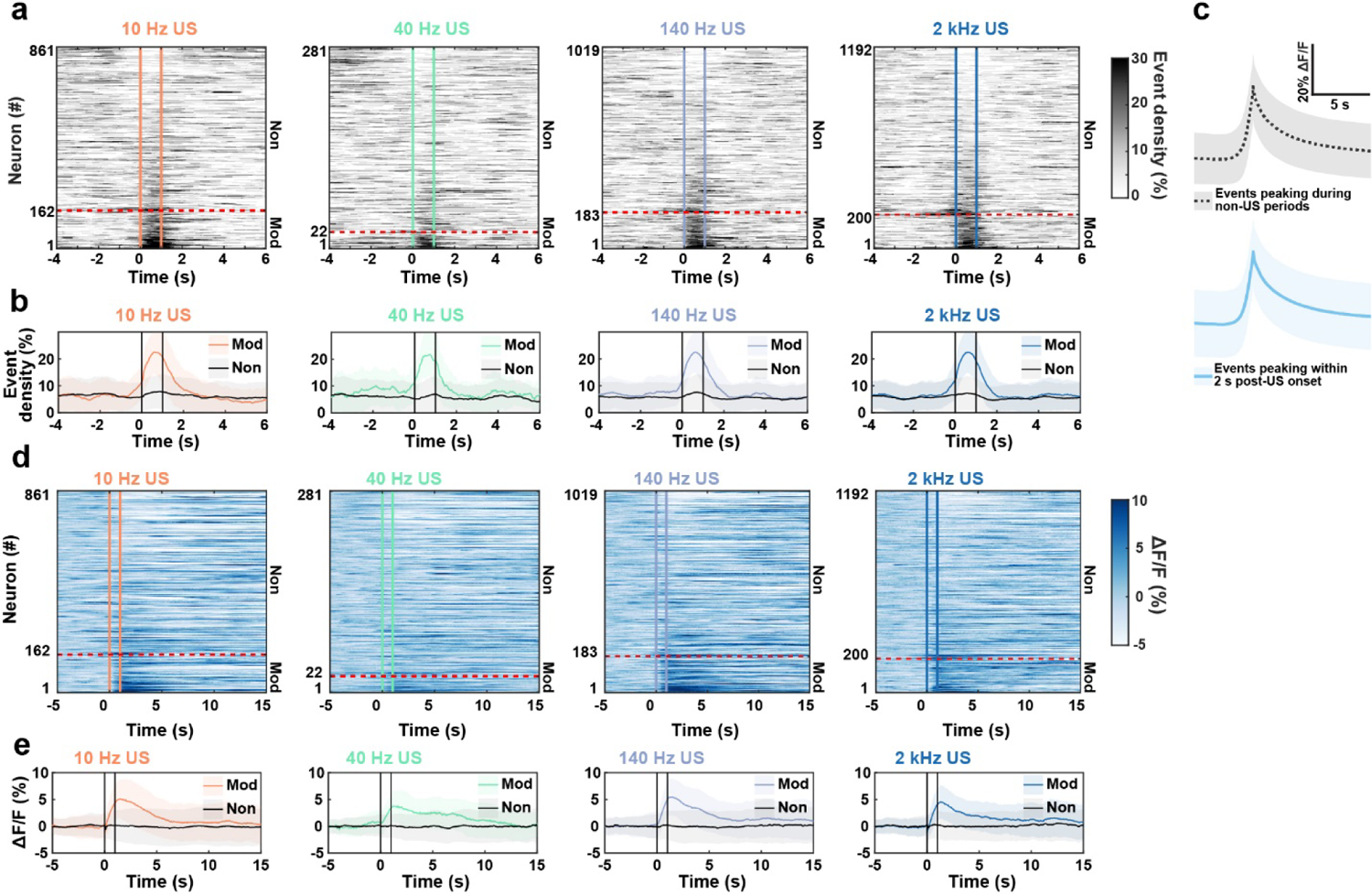
US delivered at 10 Hz, 40 Hz, and 140 Hz PRFs reliably increased the rates of calcium events associated with natural neuronal spiking. **(a)** Population average event density changes across all neurons before, during, and after US pulsed at various PRFs. Vertical colored lines represent US onset and offset. Neurons were sorted by event density difference during 1 second after versus before US onset. Neurons classified as activated by US are below the red dashed line. (Fisher’s test comparing the relative proportions of modulated neurons in stimulation vs sham conditions (**see SI 2**): 10 Hz US vs Sham p = 3.09E-15, 40 Hz US vs Sham p = 1, 140 Hz US vs Sham p = 5.63E-26, and 2 kHz US vs Sham p = 6.15E-09.) **(b)** Average modulated (orange, green, purple, and blue) and non-modulated (gray) population event density profiles. Shading represents mean event density ± SD. **(c)** Average peak-aligned calcium event waveform across events with peaks in the two seconds post-US (bottom, blue) compared to those during all other periods (top, gray). Shading represents mean ± SD. **(d)** Normalized ΔF/F heatmaps representing average calcium fluorescence activity during US. Vertical colored lines represent US onset and offset. Neurons were sorted by event density as in **2a**. Neurons classified as activated are below the red dashed line. **(e)** US-modulated and non-modulated population ΔF/F profiles. Colored lines represent modulated population profiles and gray lines represent non-modulated profiles. Shaded lines correspond to mean ΔF/F ± SD.

As a control, we recorded sham neural responses by omitting the ultrasound gel between the transducer and the mouse’s head, preventing acoustic wave propagation into the brain. Using the same statistical procedure comparing the sham-evoked response to the shuffled baseline distribution, we identified neurons activated during sham stimulation. Compiling all parameters, out of the 1915 sham neurons recorded, only 87 (4.54%) significantly increased calcium event density during sham US (**SI. 2,** Supplemental **Table 8**). The proportion of neurons modulated under the sham condition was significantly lower than the proportion modulated by US pulsed at 10 Hz, 140 Hz, and 2 kHz (Fig. 3a**, SI 2,** Supplemental **Table 7-8**, Fisher’s test, 10 Hz US vs Sham: p = 3.09E-15, 140 Hz US vs Sham: p = 5.63E-26, 2 kHz US vs Sham: p = 6.15E-09). However, 40 Hz US did not have a significant effect on modulation percentage compared to sham, likely due to smaller sample size (Fig. 3a**, SI 2,** Fisher’s test, 40 Hz US vs Sham: p > 0.05). Together, these results demonstrate that transcranial US produced large amplitude and transient cellular calcium events consistent with those occurring during natural neuron spiking.

### Transient activation of individual neurons during US does not produce long-lasting changes in network functional connectivity

The observed neural activity change in individual neurons, whether directly or indirectly responsive to US, is nonetheless associated with network changes, as activated neurons influence their downstream neurons via synaptic connections in the brain. To estimate functional connectivity, we calculated the asymmetric correlation coefficient (ACC) using the calcium event traces of each neuron pair (see **Methods**) (Fig. 4a, c, d). We first examined ACC between pairs of US-activated neurons to understand whether functional connectivity contributes to their US-sensitivity. Interestingly, pairs of US-modulated neurons had similar ACC as other cell pairs over the entire recording duration (Fig. 4b, Kruskal-Wallis test, p = 0.79), suggesting that US-sensitive neurons are not particularly functionally connected under the awake head-fixed behavioral condition.

**Figure 4.**
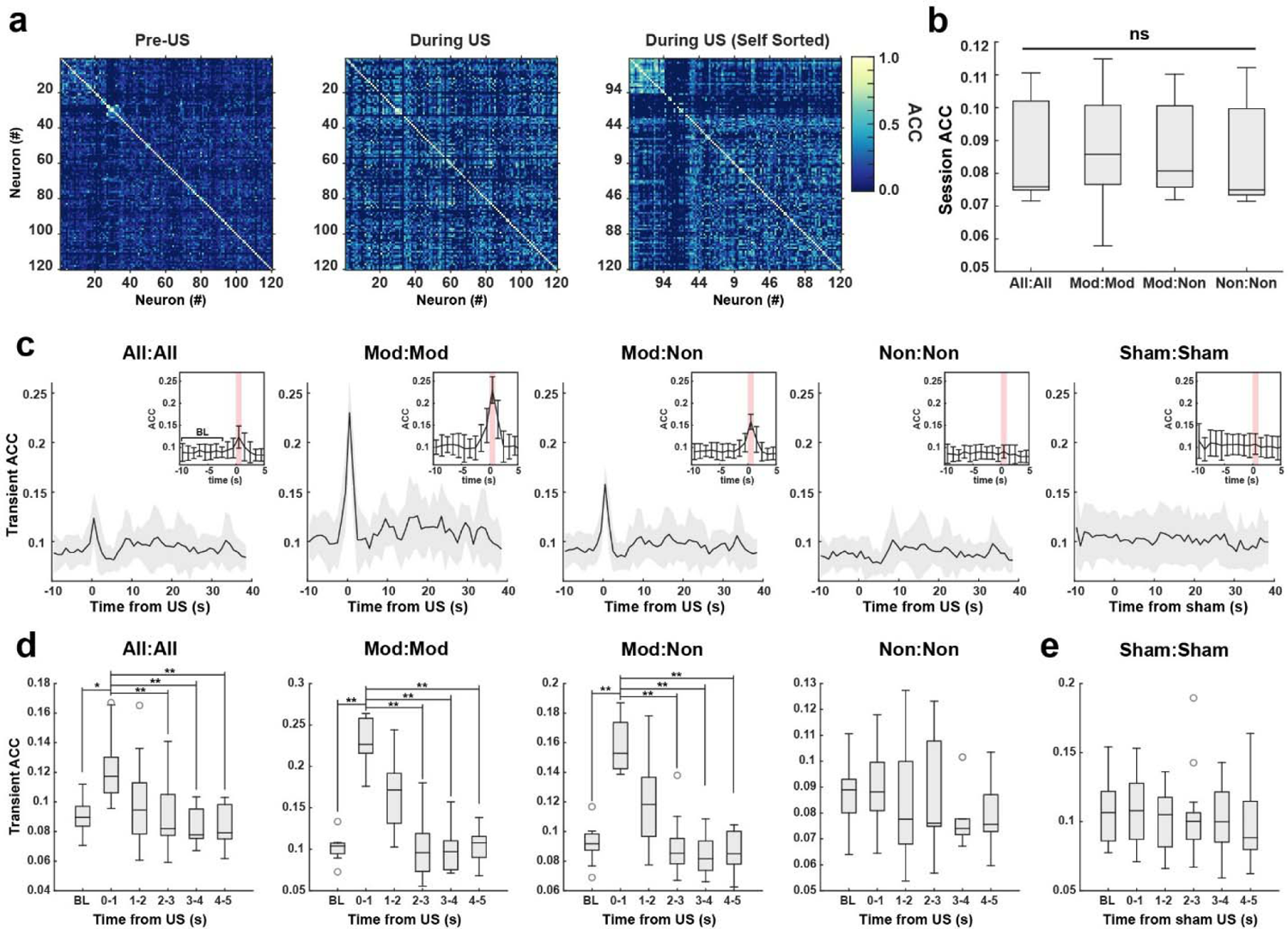
US-evoked network connectivity change is restricted to stimulation period. (**a**). Example session heatmap of ACC for cell pairs before US stimulation onset (pre-US) and during US at 10 Hz (during US). Left: ACC values were sorted with bispectral clustering to visualize subpopulations of more correlated cells. Middle: using the same sorting scheme as baseline, showing the loss of correlated clusters. Right: Resorted map during US stimulation period, showing a new cluster of correlated cells. (**b**). Session-wise averaged ACC throughout the full recording duration, for all cell pairs (“All:All”), modulated cell pairs (“Mod:Mod”), modulated and non-modulated cell pairs (“Mod:Non”), and non-modulated cell pairs (“Non:Non”). There was no significant difference for any cell pair group (Kruskal-Wallis, H(47) = 1.04, p = 0.79). (**c**) Time-course of ACC changes (“Transient ACC”) over each second for each group of cell pairs. Traces are plotted as session-wise mean ± SD. Inset: bar plot with mean ACC ± SD used for statistical comparison. The first eight seconds of the trial-averaged sessions were averaged for statistical comparison and denoted as baseline “BL”. (**d,e**) Statistical comparison of transient ACC values during the first eight seconds (“BL”) and post-US (**d**) or post sham-US (**e**). (**d**) During US (0-1 s), transient ACC was significantly higher than BL for All:All, Mod:Mod, and Mod:Non groups (All:All Friedman test, H(59) = 25.31, p = 1.21e-4, Nemenyi rank difference of US vs all other timepoints > CD for α = 0.05, Mod:Mod Friedman test, H(59) = 33.26, p = 3.35e-6, Nemenyi rank difference of US vs all other timepoints > CD for α = 0.01, and Mod:Non Friedman test, H(59) = 28.91, p = 2.41e-5, Nemenyi rank difference of US vs all other timepoints > CD for α = 0.01). There was no significant difference across timepoints for Non:Non or (**e**) sham cell pairs (Friedman test, p > 0.05).

As US transiently increased calcium event rate in modulated neurons, we next evaluated the connectivity strength between pairs of modulated neurons (Mod:Mod), between a modulated and a non-modulated neurons (Mod:Non), and between pairs of non-modulated neurons (Non:Non). As expected, ACC between neuron pairs containing a modulated neuron (Mod:Mod and Mod:Non) transiently increased during US stimulation (0-1 seconds after onset), whereas ACC between two non-modulated neurons (Non:Non) did not change (Fig. 4c, d, Friedman test, p < 0.001 for All:All, Mod:Mod, Mod:Non, and p > 0.05 for Non:Non and Sham:Sham). Across all subsets of cell pairs, increase in ACC dropped immediately at US offset and returned to baseline within a second of US offset (Fig. 4d, post-hoc Nemenyi test, BL vs 1-2, 2-3, 3-4, and 4-5 s, non-significant for all cell pair subsets). Although we detected a significant ACC increase in all recorded neuron pairs during US stimulation, a similar increase was not present during sham stimulation (Fig. 4e). Further, US-modulated neurons demonstrated no spatial clustering (**SI 3a-c**). Thus, US induces a transient increase in network synchrony largely restricted to the stimulation period with minimal sustained effects.

### Examining the differential effect of US pulsed at 10 Hz, 40 Hz and 140 Hz on parvalbumin-positive (PV) and parvalbumin-negative (non-PV) predominantly excitatory neurons

After observing reliable activation of neurons with transcranial US across PRFs, we sought to determine whether US pulsed at different PRFs produces distinct effects on the same neurons. We injected AAV9-syn-jGCaMP7f in the motor cortex of transgenic mice that selectively express the fluorophore tdTomato in PV interneurons and then placed a glass imaging window above the pia for optical access of labelled neurons (Fig. 5a). This experimental preparation allowed us to simultaneously analyze US-evoked response in PV interneurons (expressing tdTomato) and non-PV neurons (tdTomato negative, Fig. 5b). Using this experimental configuration, we performed 21 recording sessions in 6 mice and imaged a total of 2212 GCaMP7f positive neurons, including 269 (12.16%) PV cells, and 1943 (87.84%) non-PV cells. The non-PV cell population is dominated by excitatory neurons as 80-90% of neurons in the superficial motor cortex are excitatory pyramidal cells. ^56–58^

**Figure 5.**
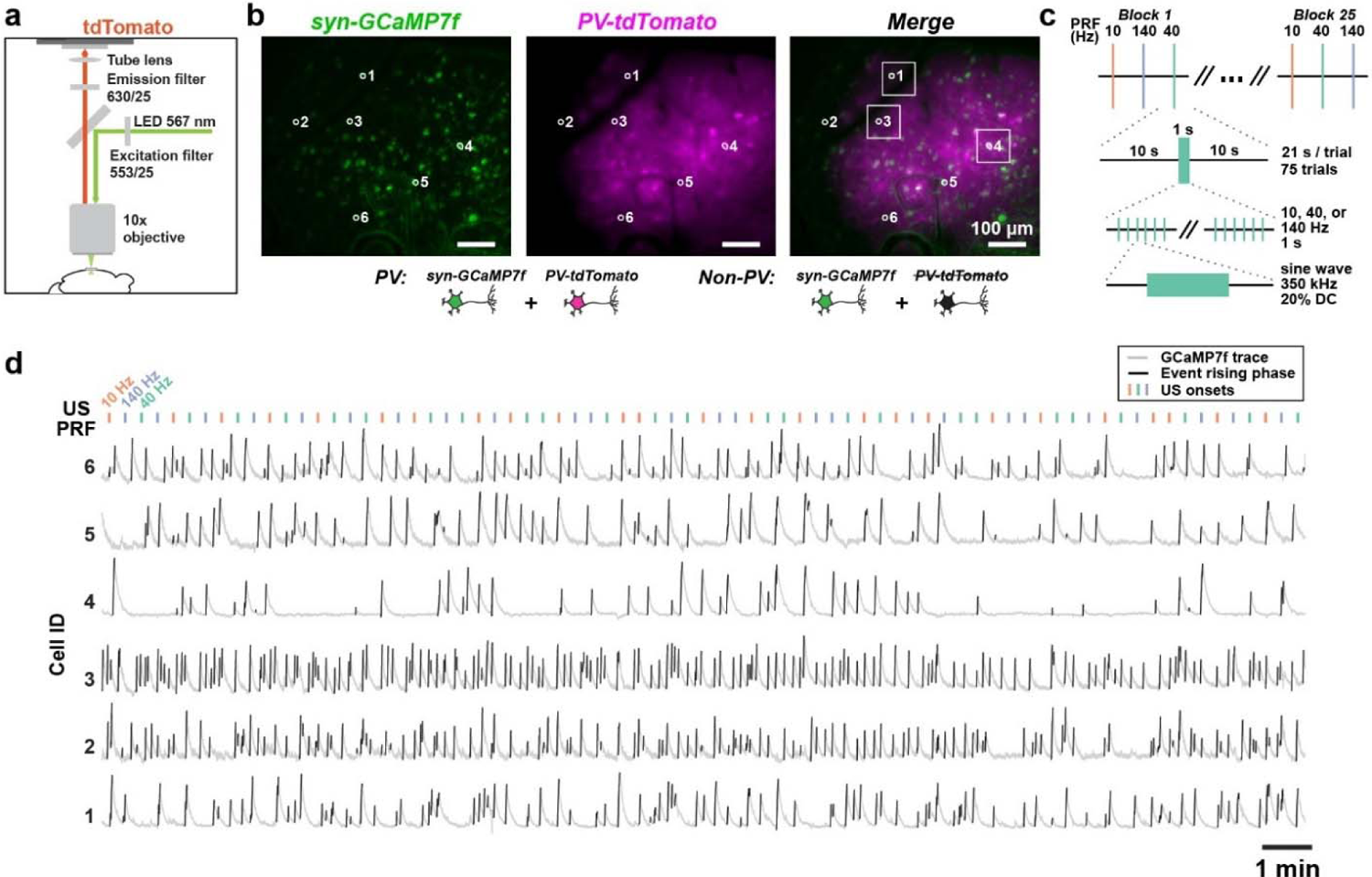
Direct comparison of the effect of different US PRF on the same neurons with or without PV expression. **(a)** Setup for imaging tdTomato (tdT) expressing PV interneurons. tdT expression was visualized using 567 nm LED excitation and 630/25 nm emissions. GCaMP7f were imaged with 460nm LED excitation and 536/40 nm emission as illustrated in Fig. 1a. **(b)** Left: An example GCaMP7f field of view, shown as fluorescence max-min projection image, with example individual neurons circled in white. Middle: Corresponding image of tdT for the same field of view. Right: Merged GCaMP7f and tdT images. PV neurons had colocalized tdT and GCaMP7f fluorescence. Neuron 4 was classified as PV, while example neurons 1, 2, 3, 5, and 6 were classified as non-PV. Boxed example neurons 1, 3, and 4 were modulated by US. All scale bars = 100 µm. **(c)** US-Stimulation trial structure for alternating PRF experiments. Each recording contained 25 blocks, with randomly alternating 10 Hz, 40 Hz, and 140 Hz trials within each block. Each trial consisted of 10 s pre-US, 1 s US, and 10 s post US. 350 kHz US was pulsed at PRFs of either 10 Hz, 40 Hz, or 140 Hz at 20% duty cycle. **(d)** Calcium fluorescence traces for example neurons during the entire recording session. Gray lines represent GCaMP7f fluorescence and black lines represent calcium event rising phases. Colored lines above depict US delivered at randomly alternating PRFs. Scale bar is 1 minute.

To understand how individual neurons respond to different PRFs, we compared the calcium responses evoked by US pulsed at 10 Hz, 40 Hz, and 140 Hz in the same neurons. As US-evoked GCaMP7f fluorescence returned to baseline levels within 10 seconds in our previous experiments (**Figs. 1, 2**), we shortened the experimental trial structure to allow for 75 total stimulation trials within one 26-minute recording period (Fig. 5c). With the increased statistical power, we divided the 75 trials into 25 blocks of alternating US PRFs at 10 Hz, 40 Hz, and 140 Hz. Each block consisted of three trials of randomly alternating PRFs, with 10 seconds pre-US, 1 second ultrasound stimulation, and 10 seconds recovery, for a total of 20 seconds between each US stimulation. We recorded GCaMP7f fluorescence during alternating PRF US and extracted fluorescence traces of individual neurons and identified calcium events (Fig. 5d).

### Most neurons were activated by only one PRF regardless of PV expression

Testing multiple PRFs on the same neuron allowed us to directly compare the sensitivity of each neuron to different US PRFs (Fig. 6a**-l**). To determine whether a neuron was activated by a specific ultrasound PRF, we compared the mean calcium event density during US of a given PRF (Fig. 6c, g, k) to the shuffled baseline event rate distribution (Fig. 6d, h, l). To generate the shuffled baseline distribution for alternating PRF experiments, we calculated the mean event rate from 25 randomly selected 1 second periods during the full recording excluding 5 seconds after all US onsets and repeated this procedure 1000 times. Neurons were deemed responsive to a specific PRF if the US-evoked event density was greater than the 95^th^ percentile of the shuffled baseline distribution. To avoid false positive identification, neurons with significantly elevated event density during the 1 second window before a given US period were excluded from being considered modulated by that PRF.

**Figure 6.**
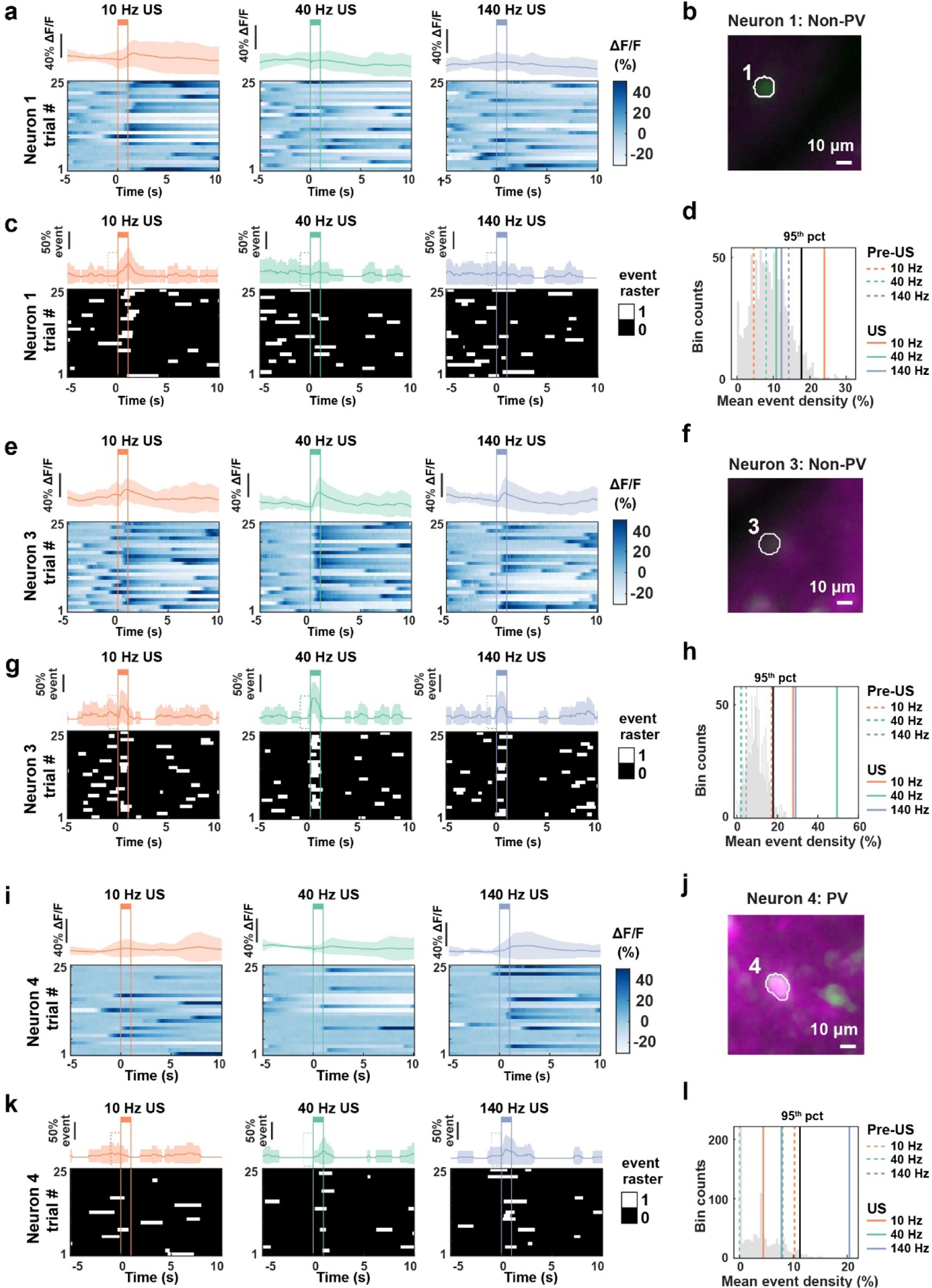
Individual neurons have different PRF preferences. (**a,e,i**) Bottom: Normalized GCaMP7f ΔF/F heatmap aligned to US onset, plotted over 25 trials for example neurons #1 (**a**), #3 (**e**), and #4 (**i**) upon US at 10 Hz (left), 40 Hz (middle) and 140 Hz (right). Top: Trial-averaged GCaMP7f ΔF/F trace. Shaded region represents mean ± SD over 25 trials. Vertical colored lines correspond to US onset and offset respectively. (**b,f,j**) Example maximum-minimum fluorescence images for the cells imaged. (**b**) non-PV neuron #1, (**f**) non-PV neuron #3, (**j**) PV neuron #4. Scale bar = 10 µm. (**c,g,k**) Bottom: Binarized event traces over 25 trials, with white corresponding to time points of event rising phases and black everywhere else. Vertical lines represent US stimulation onsets and offsets. Top: Trial-averaged event density trace. Shaded region represents mean ± SD. Dashed box prior to US depicts one second pre-US. (**d,h,l**) Illustration of statistical shuffling test used to determine neuron #1 (**d**) being modulated only by 10 Hz US, neuron #3 (**h**) being modulated by 10 Hz, 40 Hz, and 140 Hz US, and neuron #4 (**I**) being modulated only by 140 Hz US. Shuffled baseline event density is shown as a gray histogram with the 95-percentile indicated by solid black line. The observed event densities during US are shown as solid orange (10 Hz), green (40 Hz), or purple (140 Hz) lines, and the observed event densities pre-US are shown as dashed orange, green, or purple lines.

We found that both cell populations contained a sizable fraction of neurons that increased event density during US stimulation (Fig. 7a-d). Of the 269 PV neurons recorded, a similar proportion (11.2%-16.0%) were activated by US pulsed across PRFs (Fig. 7a, **SI 4a,** Supplemental **Table 9**, Chi-Square test, p = 0.26). However, the activated neurons by different PRFs were largely non-overlapping (Fig. 7b).

**Figure 7.**
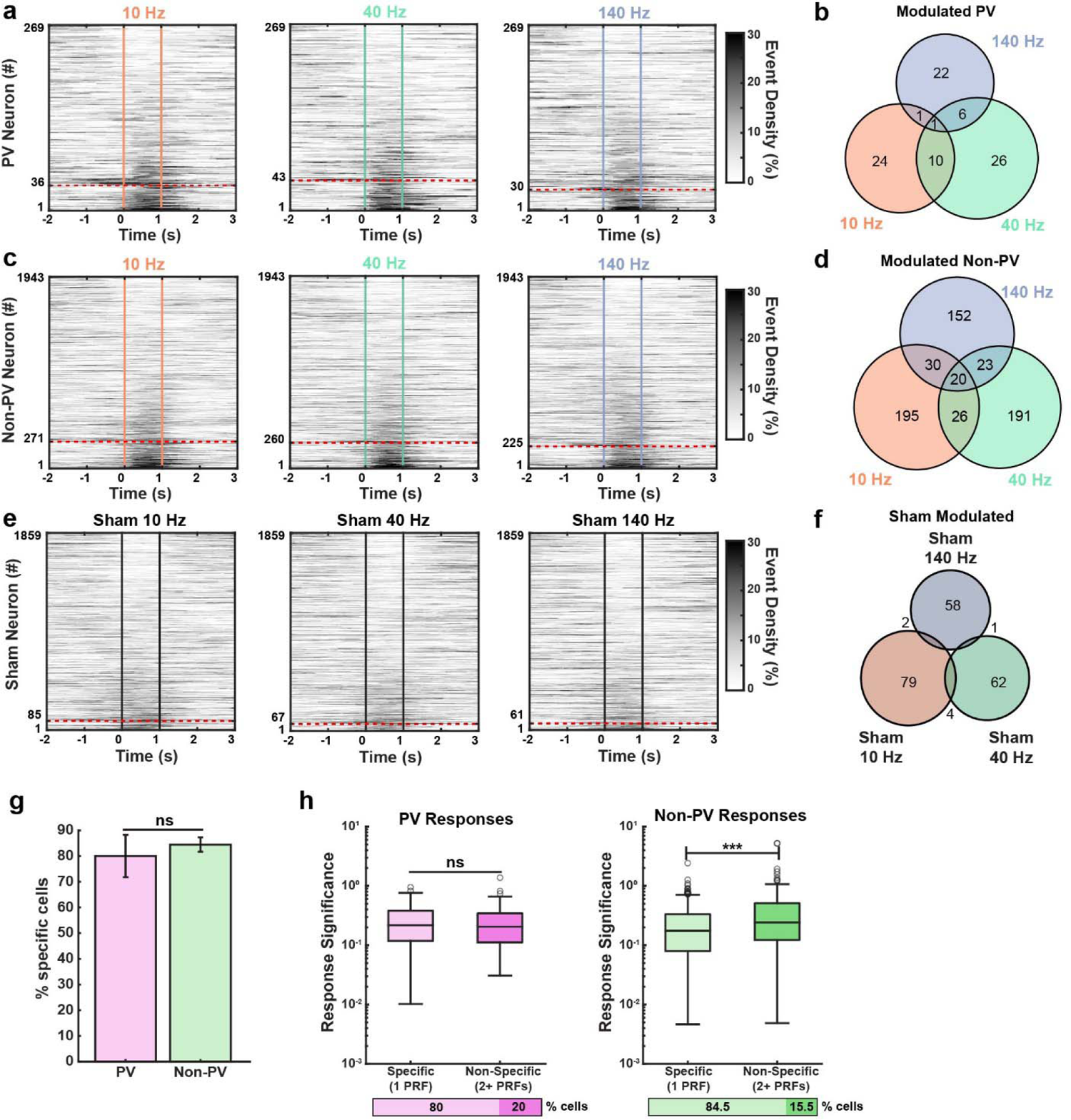
A similar proportion of largely non-overlapping cells were activated by US in both PV and non-PV populations. (**a,c,e**) Population average event density across **(a)** PV neurons, **(c)** non-PV neurons, and **(e)** sham condition. The colored lines represent US onset and offset. Neurons were sorted by event density difference during 1 second after versus before US onset. Neurons classified as activated by US are below the red dashed line. The proportion of modulated neurons in each sham condition was significantly lower than the corresponding stimulation condition (Fisher’s Test, 10 Hz US vs Sham: p = 5.27E-25, 40 Hz US vs Sham: p = 3.03E-21, and 140 Hz US vs Sham: p = 2.70e^-24^). N = 21 imaging sessions for stimulation and N = 13 imaging sessions for sham. **(b,d,f,)** Venn diagram of the cell population modulated by the specific RPFs. The number in each region corresponds to the number of cells modulated for that specific group. **(g)** Proportion of cells with responses to one PRF. Error bars represent population 95% confidence intervals. There was no significant difference between the proportion of PRF-specific cells between PV and non-PV populations (Fisher’s test, p = 0.28). **(h)** Response significance of cells responding to one PRF (“specific”) versus to two or more PRFs (“non-specific”). There was no significant difference in the response significance between non-specific and specific PV cells (Wilcoxon rank-sum test, p = 0.75, Z = 0.31). The non-specific non-PV cells had a significantly higher response significance than the specific non-PV cells (Wilcoxon rank-sum test, p = 2.27E-6, Z = −4.73). Y-axis scale is logarithmic.

Similarly, the fraction of US-activated non-PV excitatory neurons was also similar across the three PRFs tested (11.6%-14%) (Fig. 7c**, SI 4b,** Supplemental **Table 10**, Chi-Square test, p = 0.07) and largely non-overlapping (Fig. 7b). There was no difference between the fraction of activated PV and non-PV neurons for any of the PRFs (Fig. 7a, c, Fisher’s test, 10 Hz PV vs Excitatory p = 0.85, 40 Hz PV vs Excitatory p = 0.26, and 140 Hz PV vs Excitatory p = 0.92). Further, US-evoked calcium fluorescence increase was also similar between these two cell populations, both exhibiting a sharp rise followed by a gradual decay, characteristic of GCaMP7f spike-related calcium event profiles (Fig. 8a, SI 4a-b). To estimate the response latency, we averaged the calcium responses of modulated neurons across all PRFs and found a population response latency of 200 ms, corresponding to the time when the average fluorescence was greater than two standard deviations above the 10 s baseline.

**Figure 7.**
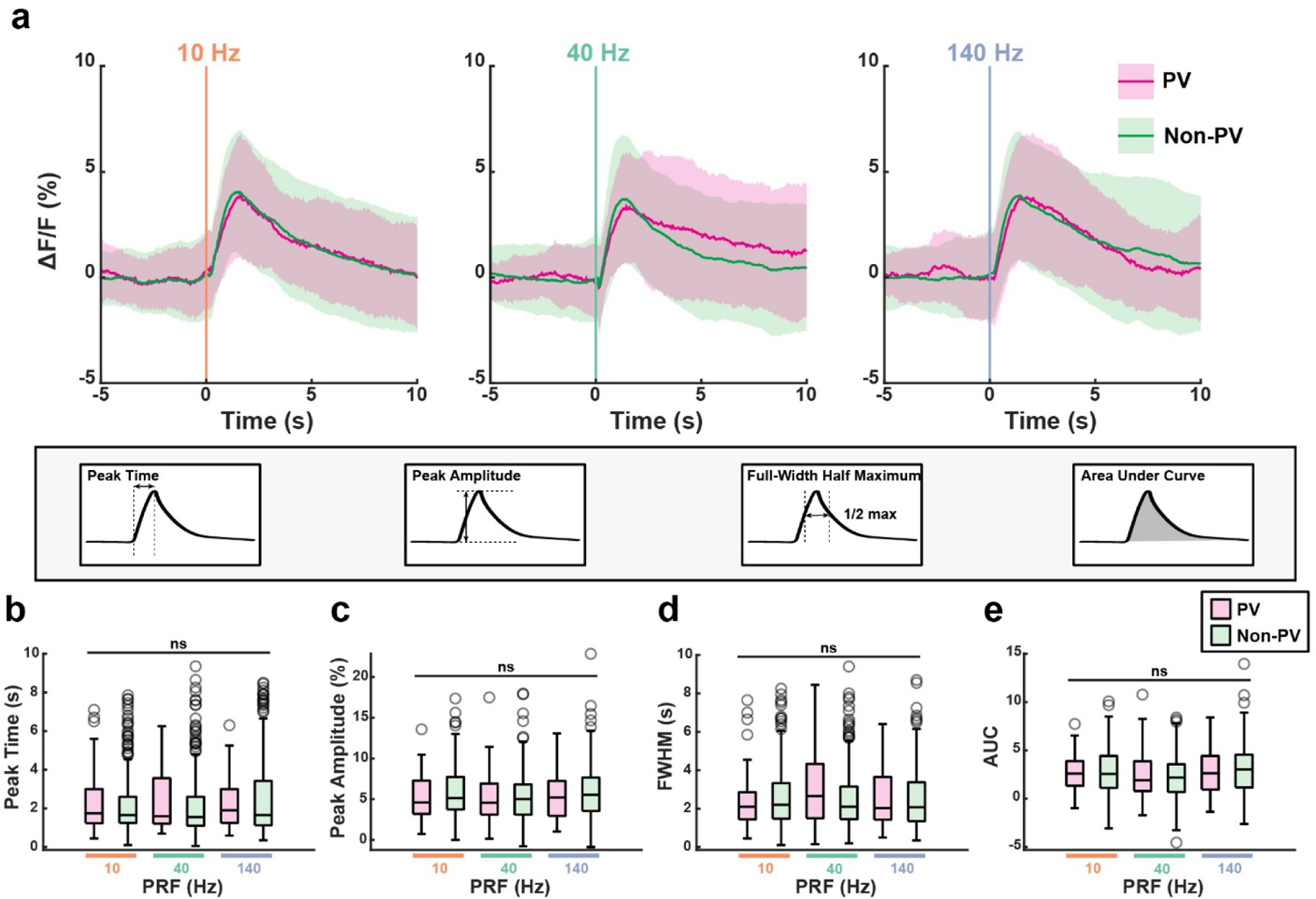
The temporal dynamics of US-evoked responses are similar across cell types and PRFs. (**a**). Average calcium fluorescence traces for 10 Hz, 40 Hz, and 140 Hz modulated PV and non-PV neurons. Shading represents mean ± SD. **(b-e)** Modulated cell’s ΔF/F calcium response profiles, characterized as time to peak (b), response amplitude (c), full width at half maximum (d) area under curve in the 5 s post-US (e). There is no difference across comparisons, except for area under curve (GLM, Deviance test, time to peak: p = 0.18; response amplitude: p=0.23, full width at half maximum: p=0.72; area under curve: p=0.045).

As a control, we employed sham stimulation by removing the ultrasound gel between the transducer and the mouse head as described previously (Fig. 7e, f). We recorded a total of 1859 neurons during sham stimulation across 13 sessions in 6 mice. Sham stimulation activated 3.3-4.6% of neurons at the PRFs tested (Fig. 7e, Supplemental **Table 11**), significantly less than that activated by US (Fig. 7a, c, b, Fisher’s test, US vs sham, p < 0.001 for 10 Hz, 40 Hz, and 140 Hz).

Interestingly, we noticed that while most neurons were responsive to one PRF (“specific” responders, examples shown in Fig. 6a-e, k-o), a small fraction responded to multiple PRFs (“non-specific” responders, examples in Fig. 6f-j). Of the 269 PV cells recorded, 72 (26.76%) were specific responders, and 18 (6.7%) were non-specific responders (Fig. 7b). Of the 1943 non-PV excitatory cells recorded, 538 (27.69%) were specific responders and 99 (5.1%) were non-specific responders (Fig. 7d). There was no difference between the fraction of specific and non-specific responders between the two cell groups (Fig. 7g, Fisher’s Test, PV vs non-PV, p = 0.28). The observation that a large fraction of neurons are specific responders, preferentially activated by only one PRF, suggests that neurons have a strong frequency response bias, potentially due to variations in mechanosensitive channel properties and their heterogeneous expression across individual neurons.

Next, we evaluated the response significance between the specific versus non-specific responders, by calculating the significance of US-evoked response defined as the difference between the observed rate minus the 95^th^ percentile of the corresponding baseline shuffled rate distribution divided by the 95^th^ percentile of the shuffled rate distribution. We found no difference between specific versus non-specific responders within the PV populations (Fig. 7h (left), Wilcoxon rank-sum test, p = 0.75). In contrast, for the non-PV population, non-specific responders had significantly higher significance than specific responders (Fig. 7h (right), Wilcoxon rank-sum test, p = 2.27E-06). As non-PV cell population includes both pyramidal cells and other inhibitory interneurons of various subtypes (Fig. 1b), the heterogeneity of US-evoked responses across non-PV cells is consistent with the idea that the PV population corresponds to a more homogeneous group of neurons than non-PV populations. The higher response significance of the non-specific responders in the non-PV population demonstrates that a subpopulation of non-PV neurons has higher sensitivity to US across PRFs, consistent with the observation that excitatory neurons have generally higher mechanosensitive ion channel expression than GABAergic neurons (Fig. 1b, f, g, SI Fig. 1).

### US-evoked calcium response profiles are similar in PV and non-PV populations across all PRFs tested

A few previous studies described small but nonetheless significantly different population responses to US between PV cells and non-PV cells ^7,16^. At the individual neuron level, we detected a similar proportion of activated cells between PV and non-PV populations. Thus, we further compared the population responses profiles between PV and non-PV excitatory neurons (Fig. 8a). We found no difference in US-evoked population calcium response peak timing, peak amplitude, and full-width half maximum between the two cell populations (Fig. 8b-d, GLM, deviance test p > 0.05). Although the area under curve (AUC) model was slightly different than the GLM intercept only model (deviance test, p = 0.0488), none of the model coefficients, including the interaction term, were significant, indicating that the model only weakly predicts the experimental values (Figure 8e, GLM coefficients p > 0.05).

There were also no major differences between the transient change in correlation coefficients among PV cell pairs, non-PV cell pairs, and PV:non-PV cell pairs (**SI 5 a-f**). As with the population of synapsin-GCaMP7f recorded previously (Fig. 4e), we found US transiently increased the correlations between cell pairs (**SI 5 b, d, f,** Friedman’s test, p < 0.05 with post-hoc Nemenyi test, BL vs US, p < 0.05 for PV, non-PV, and PV:non-PV cell pairs). After the transient increase in correlation coefficients, there was a small but significant decrease in correlation between the pairs of non-modulated non-PV cell pairs and PV:non-PV cell pairs (**SI 5d, f,** Friedman’s test p < 0.05 with post-hoc Nemenyi test BL vs 3-4 s rank difference > CD). Although PV:PV cell pairs had a similar dip in correlation, the effect was not statistically significant (**SI 5b**). Overall, correlations between cell pairs followed similar courses over experimental timepoints. Thus, the US-evoked calcium responses exhibit similar profiles between activated PV and non-PV populations. Taken together, cortical PV neurons and non-PV predominantly excitatory cells responded similarly to US stimulation at 10 Hz, 40 Hz, and 140 Hz, with a subset of each cell type preferentially responding to specific PRFs. These results provide direct experimental evidence that heterogeneity among individual cells, rather than systemic variation between cell types, dominates the effect of ultrasound neuromodulation.

## Discussion

While the underlying mechanisms behind US’s cellular effect are unclear, mechanosensitive channels are likely vital in translating US-mediated cellular effects to downstream signaling pathways^14,16,19,24,59^. We first analyzed the expression patterns of major mechanosensitive channels using single cell sequencing data obtained from the human and mice brain available at the Allen Brain Institute. Intriguingly, even though several mechanosensitive channels exhibit higher expression in excitatory neurons than inhibitory neurons, expression variation between individual neurons appears much more prominent than between cell types. This suggests that US-mediated effect is likely more heterogenous between individual cells than canonical cell types.

Next, we analyzed US-evoked responses in thousands of individual cortical neurons, with and without parvalbumin expression, using cellular GCaMP7f calcium imaging in awake mice, while pulsing 0.35 MHz US transcranially. Since the speed of neuronal activity is generally limited to a couple hundred hertz and many neurons have intrinsic biophysical properties rendering them more sensitive to certain frequency ranges, we pulsed US at the physiologically relevant frequencies of 10 Hz, 40 Hz, and 140 Hz and also included a PRF of 2 kHz. We found that all PRFs reliably evoked intracellular calcium events with similar characteristics to those naturally occurring during neuronal spiking. The calcium event activity increase across individual neurons is time locked to stimulation and accompanied by a transient increase in network synchrony. Comparing the effect of 10 Hz, 40 Hz and 140 Hz US on the same neurons, we found that most neurons were preferentially activated by a single PRF. On the population level, evoked responses were similar between the three physiological PRFs tested regardless of parvalbumin expression, suggesting that intracellular calcium changes are insensitive to variations in sonication frequencies and cell types. Together, these results highlight the consideration of PRFs in ultrasound neuromodulation and provide evidence that combining PRFs can recruit different subsets of neurons.

Most previous *in vivo* calcium imaging studies recorded neuronal responses to ultrasound at the population level without single cell resolution. In contrast, we here assessed the cellular effect across individual neurons and discovered that many neurons were preferentially activated by one PRF but not the other two. Although there is a potential for auditory and somatosensory confounds resulting from a startle-reflex at stimulation onset ^60,61^, the PRF discrimination of individual neurons cannot be explained by the broadband frequency components of the square-wave pulse onset underlying all PRFs used in our study. Further, as indirect cochlear acoustic activation is unlikely to selectively recruit motor cortex neurons at specific pulsing frequencies, the selective effects of PRF on individual neurons suggest that the observed activation is not due to indirect auditory or sensory pathway activation. While indirect cochlear activation recruited motor areas with a latency of several hundred milliseconds^60^, the population averaged response across all modulated neurons in this study had a 200 ms latency, which again cannot be fully explained by indirect auditory effects. Finally, our results are consistent with the direct neuronal effects of US on individual neurons in the hippocampal CA1 in awake mice with sub-50 ms activation latency^35^, and in eleven deafened mice^62^, *in vitro* neuron cultures^19,63^, and simple nervous systems^31^.

US activation of individual neurons involves direct engagement of mechanosensitive cellular elements, which subsequently induces intracellular cascades, recruiting voltage gated sodium and potassium channels and altering neural activity ^14,16,19,24,59^. The observed intracellular calcium increases upon US stimulation are thus expected to involve cellular signaling dependent on both plasma membrane biophysical properties (i.e., mechanosensitive ion channels) and cellular microenvironment (e.g., cell position within ultrasound wave, local chemical gradients). Our in-depth analysis of the Allen Brain Institute’s datasets revealed that many mechanosensitive channels are broadly expressed in the brains in mice and humans. Individual channels’ expression levels vary widely between brain regions. Further, the variation of channel expression is more striking among individual cells, even within a given transcriptomically defined cell type. The heterogeneity of mechanosensitive protein expression could thus result in varying US sensitivity across brain regions, cell types, and more prominently among individual cells. Consistent with this idea, we found that US sensitivity was highly heterogeneous across individual neurons, and there were no major differences between PV-positive interneurons and PV-negative predominantly excitatory neurons, suggesting that US effect is dominated by cellular signaling variations among individual cells.

Both *in vivo* and *in vitro* US neuromodulation studies implementing a wide array of parameters have demonstrated that US mediated neural responses depend on a variety of factors including brain regions ^25,35^ and cell types ^34,37^. Interestingly, US delivered at 10 Hz, 40 Hz, and 140 Hz activated comparable proportions of neurons with similar response profiles, even though many neurons were preferentially activated by only one PRF. These results support the idea that physiologically relevant PRFs differentially impact the coupling of mechano-sensing elements and downstream cellular signaling to mediate ultrasound neuromodulation. Remarkably, we did not notice any difference in US-mediated effect between PV and non-PV populations at 10 Hz, 40 Hz, or 140 Hz PRFs. Even though PV interneurons are broadly linked to 40 Hz gamma rhythms, we detected no difference in their evoked calcium responses compared to non-PV cells across PRFs, including 40 Hz.

Some recent studies reported slight differences in US-evoked responses between fast-spiking PV cells and excitatory neurons^34,37^. Our current study’s estimated acoustic pressure amplitude *in situ* was ∼0.29 MPa, lower than the 0.6 MPa used in Murphy et al ^34^, potentially contributing to our insignificant differences between cell types. We also found that US-evoked calcium events exhibited similar characteristics as those naturally occurring during increased spiking in the absence of US stimulation. However, as calcium imaging is slow, we may have missed the subtle differences in US-evoked spiking activity captured by extracellular recordings of putative PV positive fast spiking cells and putative excitatory broad spiking cells ^37^.

Previous *in vivo* studies detected diverse neural responses to ultrasound across multiple brain regions ^25,35,64^. However, none directly compared the neuronal response to multiple parameters at the single cell level across different cell types. We found that most neurons were preferentially activated by a single PRF. Combining PRFs, such as the three tested here, could thus enhance the efficacy of US neuromodulation by recruiting different subsets of neurons in clinical applications. With future investigation on the effect of adjusting or combining PRFs to target desired neuronal subgroups, it may be possible to design personalized medicine opportunities. While the majority of PV and non-PV neurons selectively responded to a single PRF, some responded non-selectively to two or more. Heterogenous expression of mechanosensitive channels with varying resonant frequencies may contribute to PRF dependent activation between different neurons within the same cell type. It is also highly plausible that non-specific responders have cellular environments that are more sensitive to US, allowing them to be responsive to multiple PRFs.

Despite consistent calcium dynamics observed in both responding cell types, only the non-PV population exhibited variations in response significance when comparing selective and non-selective responders. The non-PV cells that responded to multiple PRFs exhibited significantly greater responses than those that selectively responded to single PRFs. Conversely, the effect size of responding PV cells remained consistent regardless of PRF selectivity. As the non-PV cell population comprises mainly pyramidal neurons, heightened response significance in non-PV cells is consistent with higher mechanosensitive channel expression observed in our single cell gene expression analysis.

Numerous factors influence neuronal excitability including membrane potential, network activity, and channel distribution. Using statistical resampling tests, we identified about 10-20% of neurons activated by US by a single PRF. However, US evoked responses within the same neurons exhibited large variations across trials, confirming US’s weak modulatory effect. One potential explanation for this weak effect is that US depolarizes neurons through mechanosensitive channel activation, thus indirectly increasing spiking probability via downstream signal amplification by voltage and calcium gated channels ^19,59^. Nonetheless, the overall weak effect is consistent with the behavioral state, brain region and pulse pattern dependent effect of US broadly observed across various studies^34,35,37,64^. The underlying mechanisms of ultrasonic neuromodulation and selective sensitivity to PRFs remain unknown. Future studies looking into mechanosensitive channel sensitivity to US and its role in engaging cellular signaling pathways that govern neuronal excitability will further clarify the nature of US stimulation and frequency response bias.

## Methods

### Allen Brain Institute Gene expression analysis

Data are from Allen Institute for Brain Science, https://brain-map.org/atlases-and-data/rnaseq/human-m1-10x and https://brain-map.org/atlases-and-data/rnaseq/mouse-whole-cortex-and-hippocampus-10x. Uniform Manifold Approximation and Projection (UMAP) representation of the mouse single cell sequencing results were created with Cytosplore Viewer, https://viewer.cytosplore.org/. To normalize expression levels for individual cell comparisons, raw unique molecular identifier (UMI) counts were divided by the total UMI count for the given cell, multiplying by a scaling factor of 100,000, adding 1, and performing a log-2 transform. Expression level distribution overlap is measured with Cliff’s delta (8), with |8| < 0.147 considered negligible, 0.147 < |8| < 0.330 considered small, 0.330 < |8| < 0.474 considered medium, and |8| > 0.474 considered large.

### Animal preparation

All animal experimental procedures were approved by the Boston University Institutional Animal Care and Use (IACUC) and Biosafety Committees. When possible, mice were group housed prior to surgery and single-housed post-surgery. Enrichment was provided with Igloos or running wheels. Animal facilities were maintained around 70°F and 50% humidity and were kept on a 12-hour light/dark cycle.

We used a total of 16 adult mice, including 10 C57BL6 mice (9 female and 1 male) and 6 PV-tdTomato transgenic mice that were generated by crossing B6.Cg-*Gt(ROSA)26Sor^tm^*^14^*^(CAG-tdTomato)Hze^*/J (Jax, stock number: 007914) with B6.129P2-*Pvalb^tm1(cre)Arbr^*/J (Jax, stock number: 017320) mice (3 male and 3 female) (Jackson Laboratory, Bar Harbor, ME). Mice were 8-16 weeks old at the time of surgery.

Animal preparations are identical to those described previously ^35,65^. Briefly, under isoflurane general anesthesia, we first performed a craniotomy ∼3mm in diameter over the right motor cortex (AP: 1.75 mm, ML: 1.75 mm). We then infused a total of 0.5uL of AAV9-syn-GCaMP7f (titer 2.3×10^13^ GC/ml, Addgene 104488-AAV9, Watertown, MA) at two separate locations within the craniotomy, about 180-nm below the dura, at 80 nl/min using a World Precision Instruments NanoFil syringe (NANOFIL) with a 36-gauge bunt needle (NF36BL-2). Following injection, the injection needle was left in place for 10 minutes at each site to allow AAV diffusion. We then placed a cover glass (no. 0, OD: 3 mm, Deckgläser Cover Glasses, Warner Instruments, 64-0726 (CS-3R-0)) over the viral injection site and fixed it in place using the UV curable glue (Tetric EvoFlow (Safco, Buffalo Grove, IL). Metabond Quick Adhesive Cement System (SKU: S380 Parkell, Edgewood, NY) was then used to cover any exposed skull, followed with dental cement (5145, Stoelting, Wood Dale, IL) to fix a custom metal head-bar posterior to the motor cortex near lambda. Mice were administered preoperatively with buprenex (0.1mg/kg) or with sustained release (SR) buprenorphine (3.25mg/kg, Ethiqa XR, Fidelis Pharmaceuticals, North Brunswick Township, NJ) postoperatively to provide a minimum of 48 hours analgesia. Following surgery, mice were given a 2–3-week recovery period before recording sessions began.

### Calcium imaging and ultrasound stimulation

Mice were habituated for one week and then imaged using a custom wide-field microscope equipped with a 10x objective at NA 0.28 (10x M Plan APO-378-803-3, Mitutoyo, Takatsu-ku, Kawasaki, Kanagawa, Japan). A 460 nm LED (LZ1-00B200; LedEngin, San Jose, CA) with an excitation filter at 470/50 nm (FF01-468/553-25; Semrock, Rochester, NY) was used to excite GCaMP7f. GCaMP7f fluorescence emissions then passed through a dichroic mirror (FF493/574-Di01-25×36; Semrock, Rochester, NY) and were filtered at 536/40 nm (FF01-512/630-25; Semrock, Rochester, NY). GCaMP7f imaging was performed at 20 Hz with a sCMOS camera (Hamamatsu ORCA Fusion C14440-20UP; Hamamatsu Photonics K.K., Shizuoka, Japan).

Mice were positioned below the objective and head-fixed on a 3D printed platform, directly above a planar ultrasound transducer with 350 kHz center frequency (GS350-D13, Ultran, State College, PA). The chin of the animals was placed at the center of the ultrasound transducer, and the space between the chin and the transducer was filled with ultrasound gel (Aquasonic Clear® Ultrasound Gel 03-08_BX, Parker Laboratories, Fairfield, NJ) to ensure ultrasound propagation to the brain. Sham stimulations were performed without ultrasound gel between the chin and the transducer.

Two ultrasound stimulation protocols were used in this study. For the first protocol, we recorded long duration trials while delivering US at a single pulse repetition frequency (PRF). For these experiments, each recording session had 20 imaging trials, with each trial lasting for 50 seconds containing a 10 second period before US followed by 1 second of ultrasound stimulation and 39 seconds recovery, for a total inter-stimulation interval of 49 seconds. Ultrasound PRF of 10 Hz, 40 Hz, or 140 Hz at 20% DC or 2 kHz at 42% DC was used. There was a 0.5 second pause between each trial, and GCaMP7f fluorescence traces were interpolated to account for missing data points between trials. For the second protocol, we recorded shorter duration trials with PRF alternating between 10 Hz, 40 Hz, and 140 Hz at 20% DC during each session. Specifically, each recording session consisted of 25 blocks, with each block containing a randomly permuted PRF combination of 3 trials at 10 Hz, 40 Hz, and 140 Hz respectively. Thus, each recording contained 75 trials, with 25 trials per PRF. Each trial began with a 10 second period without US followed by 1 second of ultrasound stimulation and 10 seconds post-stimulation. There was a 0.5 second pause between each trial to maintain signal alignment, and the resulting GCaMP7f fluorescence traces were interpolated to account for missing data points.

Ultrasound parameters including PRF, stimulation duration, and ultrasound onset time were programmed in MATLAB (MATLAB r2021b, MathWorks, Natick, MA), which sent TTL pulses through a NI-DAQ multifunction I/O system (PCIe-6321, National Instruments, Austin, TX) to trigger camera acquisition and ultrasound pulses. For ultrasound stimulation, MATLAB programmed TTL pulses triggered a function generator (33220A, Keysight, Santa Rosa, CA) to generate a 350 kHz sine wave at the desired repetition frequency and duty cycle. The function generator output was connected to an amplifier (AG1006, T&C Power Conversion, Rochester, NY) to amplify the voltage signal at 350 kHz. The amplified voltage signal was sent to the ultrasound transducer, which converted the voltage to an acoustic wave. OpenEphys (Open Ephys Acquisition Board v2.4, OpenEphys, Atlanta, GA) acquired individual imaging frame timestamps from the camera and all TTL triggers from the NI-DAQ to allow for offline alignment between ultrasound stimulation onsets and calcium imaging frames.

### Hydrophone calibration and intensity calculations

Acoustic intensity measurements were performed with a needle hydrophone calibrated between 0.25-1 MHz (HNR-1000, Onda Corporation, Sunnyvale, CA) and connected to a pre-amplifier (AH-2010, Onda Corporation). The hydrophone was positioned above the 350 kHz ultrasound transducer in degassed water. The free-field spatial peak of the ultrasound beam was found by spatially translating the hydrophone in the x, y, and z planes while monitoring peak-peak voltage on an oscilloscope. After finding the spatial peak, the hydrophone waveform was recorded with a high-speed 14-bit digitizer card (Octave CSE8325, GaGe by VITREK, Lockport, IL). Voltage waveforms were converted to pressure with the calibrated sensitivity at 350 kHz adjusted for the preamplifier (7.234×10^-6^ V/Pa). We estimated free-field spatial-peak pulse-average intensity (I_SPPA_) with the following equation: 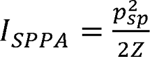, where *p_sp_* is the spatial peak pressure amplitude and the acoustic impedance of water is Z::: 1.5 × 10^6^ Rayls. Spatial-peak, temporal average intensity was then calculated by multiplying I_SPPA_ by the duty cycle of stimulation.

### Ultrasound acoustic pressure simulation

Acoustic pressure was simulated with the k-Wave acoustic modeling toolbox in MATLAB as described previously in Tseng et al 2021^35^. Briefly, acoustic pressure was simulated within a three-dimensional representation of a C57BL/6 mouse skull (adapted from Chan et al 2007)^54^. The skull model was modified to include a craniotomy and a glass coverslip above the motor cortex, and two air pockets were added in the mouth to simulate acoustic propagation from below the chin. Both soft tissue and bone were considered homogeneous with bulk acoustic properties, and the mouse head model was surrounded by air at 50% humidity and 25°C. As in the experiments, the mouse head was coupled to the acoustic source via ultrasound gel, which was modeled as water. The simulation’s acoustic source was a 12.36 mm diameter single element planar ultrasound transducer. The k-Wave computational grid was discretized at 0.12 mm and the transmitted ultrasound was a 25-cycle burst at 350 kHz with peak pressure of 522 kPa, as estimated with hydrophone measurements. The k-Wave simulation estimated the maximum instantaneous acoustic pressure and intensity throughout the brain during ultrasound stimulation. To validate simulation results, we measured the acoustic pressure in a mouse skull filled with ultrasound gel with a hydrophone.

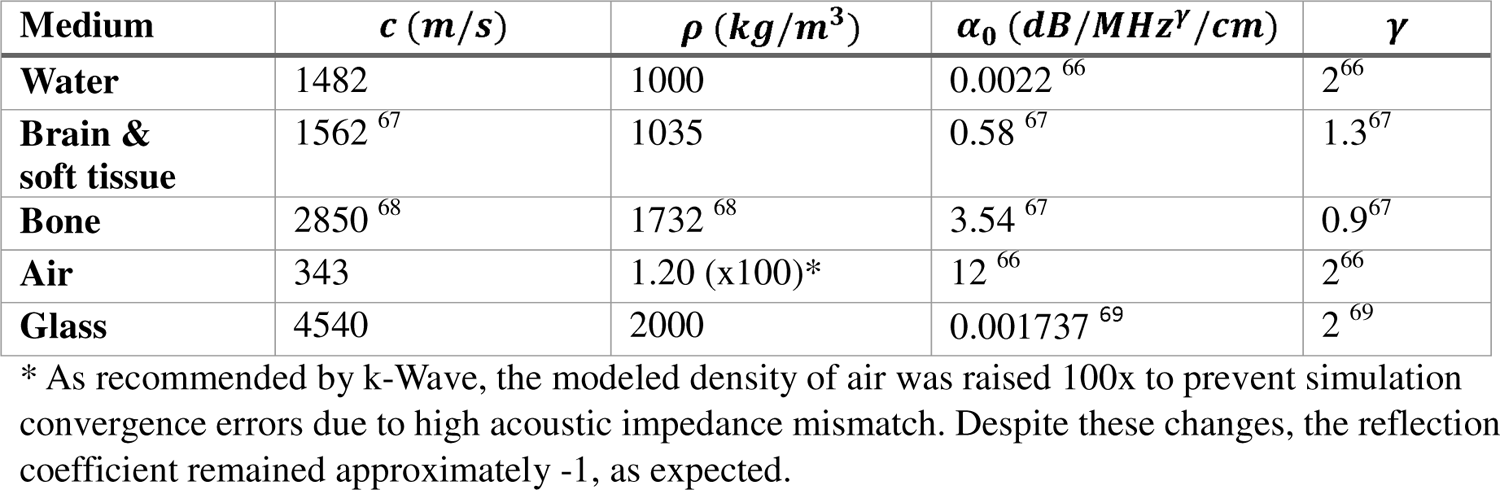

| Medium | $c$ (m/s) | $\rho$ (kg/m<sup>3</sup>) | $\alpha_0$ (dB/MHz<sup>γ</sup>/cm) | $\gamma$ |
| --- | --- | --- | --- | --- |
| Water | 1482 | 1000 | 0.0022<sup>66</sup> | 2<sup>66</sup> |
| Brain & soft tissue | 1562<sup>67</sup> | 1035 | 0.58<sup>67</sup> | 1.3<sup>67</sup> |
| Bone | 2850<sup>68</sup> | 1732<sup>68</sup> | 3.54<sup>67</sup> | 0.9<sup>67</sup> |
| Air | 343 | 1.20 (x100)* | 12<sup>66</sup> | 2<sup>66</sup> |
| Glass | 4540 | 2000 | 0.001737<sup>69</sup> | 2<sup>69</sup> |

\* As recommended by k-Wave, the modeled density of air was raised 100x to prevent simulation convergence errors due to high acoustic impedance mismatch. Despite these changes, the reflection coefficient remained approximately -1, as expected.

### Calcium imaging ROI segmentation and GCaMP7f trace extraction

Calcium videos were first motion corrected as described in Tseng et al 2021^35^, and then regions of interest (ROIs) corresponding to individual neurons were automatically segmented using a custom deep learning network followed by manual inspection to remove ROIs that do not exhibit neuronal morphologies. Additional ROIs missed by the algorithm were added during manual inspection in MATLAB. The GCaMP7f trace for each neuron was calculated as the mean intensity of all pixels within a ROI minus the mean pixel intensity of the surrounding ‘donut’ masks (inner radius 15 pixels from ROI centroid and outer radius 50 pixels from ROI centroid) to remove potential local background activity.

Pixels within ROIs were excluded from the background donut mask. GCaMP7f traces were interpolated to 20 Hz using shape-preserving piecewise cubic interpolation and linearly detrended to remove photobleaching effects. Finally, GCaMP7f traces were normalized between 0 and 1 by subtracting the minimum trace value and dividing by the fluorescence intensity range. To ensure similar values in the 10 s pre-US period throughout trials, the final ΔF/F value was calculated by subtracting the average value of the10 s pre-US period for each trial from that trial’s trace.

### Parvalbumin (PV) positive interneuron detection

To identify PV cells among the recorded GCaMP7f expressing neurons, we captured tdTomato fluorescence at 100 ms exposure, immediately prior to GCaMP7f recording. From the tdTomato fluorescence image, we marked matching vascular landmarks in the tdTomato images and GCaMP7f images and warped the tdTomato image to match the GCaMP7f vascular landmarks (MATLAB estimateGeometricTransform and imwarp functions). Next, we identified tdTomato positive cells (tdTomato ROIs) using the neuron segmentation method detailed above to identify GCaMP7f positive neurons (GCaMP7f ROIs). We then identified PV cells as GCaMP7f ROIs with >50% overlap with tdTomato ROIs.

### Calcium event detection

GCaMP7f calcium events were detected using the frequency spectral profile of their sharp rise in fluorescence, as described previously ^65^. Briefly, each GCaMP7f trace was smoothed with a 1 second sliding window (MATLAB movmean). Next, the continuous multi-taper time-frequency spectrum was found for each smoothed GCaMP7f trace (Chronux Toolbox in MATLAB, http://chronux.org). Potential event peaks were found as points in the frequency spectrum above the median normalized power.

Potential events with rise times greater than 100ms and amplitudes greater than 2.5x the pre-event standard deviation were classified as calcium events. Rise time was defined as the time between the minimum trace timepoint within 3 seconds before the peak and the peak timepoint. To ensure quality GCaMP7f event data, GCaMP7f traces without any events throughout the entire recording duration were removed from further analysis. Remaining traces were manually inspected to ensure calcium events exhibited fast rise and exponential decay with minimal noise.

### Identification of neurons activated by ultrasound

To determine whether a neuron was activated by ultrasound stimulation, we first binarized the calcium event traces using ones for the rising phase and zeros everywhere else. We then calculated the ultrasound evoked responses as the mean event rate over the 1 second period after US onset. US-evoked event rate in each neuron was then compared to the baseline shuffled distribution of the given neuron. We defined the baseline period for each neuron as the entire recording period excluding the 5 second periods following each US onset. To create the baseline shuffled distribution of event rates, we randomly selected 25 of 1-second-long periods during the baseline period during each iteration and calculated the mean calcium event rate. This procedure was iterated 1000 times with replacement. If a neuron’s US-evoked event rate is above 95^th^ percentile of the shuffled baseline distribution, the neuron was deemed significantly activated by ultrasound. If US-evoked event rate is within 95^th^ percentile of the baseline shuffled distribution, the neuron was deemed not modulated.

### Pair-wise asymmetric correlation analysis

The asymmetric correlation coefficient (ACC) was used to estimate neural calcium event synchrony, as it represents the proportion of co-occurring calcium events between two neurons. Each neuron’s binarized calcium event trace was compared to all other neurons with the following equation:

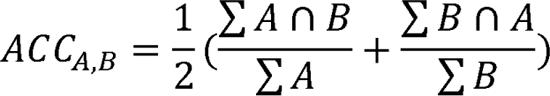

To determine if ultrasound induces any long-term changes in network synchronization, we calculated the sustained ACC using the full recording duration’s calcium event traces. Short-term changes in calcium event synchrony were estimated by calculating the transient ACC over one second bins throughout the recording. For statistical comparison of ACCs between different timepoints or cellular populations, we used session averaged ACC values.

### Ultrasound-evoked GCaMP7f response characteristics

To compare population ultrasound-evoked GCaMP7f fluorescence changes, we first computed US-evoked GCaMP7f fluorescence changes in each neuron by aligning GCaMP7f trace to US onset. Evoked peak fluorescence was defined as the maximum fluorescence value within 5 seconds of US onset. If the peak was found after 5 seconds, the neuron was excluded from further response analysis.

Evoked response amplitude was calculated as the difference between evoked peak fluorescence and the minimum fluorescence value of the GCaMP7f trace between US onset and the evoked peak. Peak timing was the time from the ultrasound onset to the peak. The full-width half-maximum (FWHM) was defined as the time GCaMP7f fluorescence remained above 50% of the evoked response amplitude. The area under the curve (AUC) was calculated with trapezoidal numerical integration during the 5 s post-US onset (trapz function in MATLAB).

### Statistical analysis

We used significance level α = 0.05 for all statistical tests. Using quantile-quantile plots and the Shapiro-Wilkes test for normality, we determined that most of our data was not normally distributed.

Accordingly, we used non-parametric statistical tests for our data analysis. When comparing relative proportions of modulated and non-modulated cells between two groups, we used Fisher’s exact test. When comparing relative proportions between three or more groups, we used a chi-square test. For comparison of medians between two groups of matched pairs, we used the Wilcoxon signed-rank test, and for comparisons between two groups of unmatched pairs we used the Wilcoxon rank-sum test. To compare medians of groups of three or more, we employed the non-parametric Kruskal-Wallis test. If a Kruskal-Wallis test was significant (p < 0.05), we ran a post-hoc multiple comparisons test with Dunn-Sidak corrections to test significance between groups. For comparisons of multiple repeated measures, we used a Friedman test with a post-hoc Nemenyi test.

For analyzing the influence of multiple predictor variables (i.e., pulse repetition frequency (PRF) and cell type) on a response variable Y (i.e., calcium response shape characteristics), we used generalized linear models (GLM, fit with function MATLAB fitglm). We used the same GLM architecture for all response variables:

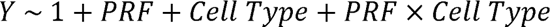

Each GLM’s fit was compared to an intercept-only model with the deviance test, and we only analyzed the GLM coefficients if the deviance test was significant. As none of the GLM coefficients were significant, we did not perform post-hoc analyses.

## Acknowledgements

XH acknowledges funding from NSF (CBET-1848029 and CIF-1955981) and NIH R01NS109794. JS acknowledges NIH T32-GM008541. EB acknowledges NSF GRFP DGE-1840990 and NIH T32-EB006359. ES acknowledges NIH T32-GM008764. The authors acknowledged computational work done on the Shared Computing Cluster in Boston University’s Research Computing Services.

## References

1. Beisteiner, R., Hallett, M. & Lozano, A. M. Ultrasound Neuromodulation as a New Brain Therapy Advanced Science 10, (2023).

2. Hu, Y.-Y. et al. Transcranial low-intensity ultrasound stimulation for treating central nervous system disorders: A promising therapeutic application. Front Neurol 14, (2023).

3. Legon, W., Bansal, P., Tyshynsky, R., Ai, L. & Mueller, J. K. Transcranial focused ultrasound neuromodulation of the human primary motor cortex. Sci Rep 8, 10007 (2018).

4. Lee, W. et al. Transcranial focused ultrasound stimulation of human primary visual cortex. Sci Rep 6, 34026 (2016).

5. Lee, W. et al. Image-Guided Transcranial Focused Ultrasound Stimulates Human Primary Somatosensory Cortex. Sci Rep 5, 8743 (2015).

6. Gibson, B. C. et al. Increased Excitability Induced in the Primary Motor Cortex by Transcranial Ultrasound Stimulation. Front Neurol 9, (2018).

7. Ai, L., Bansal, P., Mueller, J. K. & Legon, W. Effects of transcranial focused ultrasound on human primary motor cortex using 7T fMRI: a pilot study. BMC Neurosci 19, 56 (2018).

8. Mueller, J., Legon, W., Opitz, A., Sato, T. F. & Tyler, W. J. Transcranial Focused Ultrasound Modulates Intrinsic and Evoked EEG Dynamics. Brain Stimul 7, 900–908 (2014).

9. Lemaire, T., Neufeld, E., Kuster, N. & Micera, S. Understanding ultrasound neuromodulation using a computationally efficient and interpretable model of intramembrane cavitation. J Neural Eng 16, 046007 (2019).

10. Dell’Italia, J., Sanguinetti, J. L., Monti, M. M., Bystritsky, A. & Reggente, N. Current State of Potential Mechanisms Supporting Low Intensity Focused Ultrasound for Neuromodulation. Front Hum Neurosci 16, (2022).

11. Taylor, G. J. et al. Capacitive Detection of Low-Enthalpy, Higher-Order Phase Transitions in Synthetic and Natural Composition Lipid Membranes. Langmuir 33, 10016–10026 (2017).

12. Blanco-González, A., Marrink, S. J., Piñeiro, Á. & García-Fandiño, R. Molecular insights into the effects of focused ultrasound mechanotherapy on lipid bilayers: Unlocking the keys to design effective treatments. J Colloid Interface Sci 650, 1201–1210 (2023).

13. Man, V. H., Li, M. S., Wang, J., Derreumaux, P. & Nguyen, P. H. Interaction mechanism between the focused ultrasound and lipid membrane at the molecular level. J Chem Phys 150, (2019).

14. Tyler, W. J. The mechanobiology of brain function. Nat Rev Neurosci 13, 867–878 (2012).

15. Naor, O., Krupa, S. & Shoham, S. Ultrasonic neuromodulation. J Neural Eng 13, 031003 (2016).

16. Plaksin, M., Kimmel, E. & Shoham, S. Cell-Type-Selective Effects of Intramembrane Cavitation as a Unifying Theoretical Framework for Ultrasonic Neuromodulation. eNeuro 3, ENEURO.0136-15.2016 (2016).

17. Tyler, W. J. Noninvasive Neuromodulation with Ultrasound? A Continuum Mechanics Hypothesis. The Neuroscientist 17, 25–36 (2011).

18. Menz, M. D. et al. Radiation Force as a Physical Mechanism for Ultrasonic Neurostimulation of the *Ex Vivo* Retina. The Journal of Neuroscience 39, 6251–6264 (2019).

19. Yoo, S., Mittelstein, D. R., Hurt, R. C., Lacroix, J. & Shapiro, M. G. Focused ultrasound excites cortical neurons via mechanosensitive calcium accumulation and ion channel amplification. Nat Commun 13, 493 (2022).

20. Kefauver, J. M., Ward, A. B. & Patapoutian, A. Discoveries in structure and physiology of mechanically activated ion channels. Nature 587, 567–576 (2020).

21. Brohawn, S. G., Su, Z. & MacKinnon, R. Mechanosensitivity is mediated directly by the lipid membrane in TRAAK and TREK1 K ^+^ channels. Proceedings of the National Academy of Sciences 111, 3614–3619 (2014).

22. Sorum, B., Rietmeijer, R. A., Gopakumar, K., Adesnik, H. & Brohawn, S. G. Ultrasound activates mechanosensitive TRAAK K ^+^ channels through the lipid membrane. Proceedings of the National Academy of Sciences 118, (2021).

23. Zhang, M., Shan, Y., Cox, C. D. & Pei, D. A mechanical-coupling mechanism in OSCA/TMEM63 channel mechanosensitivity. Nat Commun 14, 3943 (2023).

24. Qiu, Z. et al. The Mechanosensitive Ion Channel Piezo1 Significantly Mediates In Vitro Ultrasonic Stimulation of Neurons. iScience 21, 448–457 (2019).

25. Zhu, J. et al. The mechanosensitive ion channel Piezo1 contributes to ultrasound neuromodulation. Proceedings of the National Academy of Sciences 120, (2023).

26. Yang, Y. et al. Induction of a torpor-like hypothermic and hypometabolic state in rodents by ultrasound. Nat Metab 5, 789–803 (2023).

27. Inoue, R., Jian, Z. & Kawarabayashi, Y. Mechanosensitive TRP channels in cardiovascular pathophysiology. Pharmacol Ther 123, 371–385 (2009).

28. Lau, O.-C. et al. TRPC5 channels participate in pressure-sensing in aortic baroreceptors. Nat Commun 7, 11947 (2016).

29. Starostina, I. et al. Distinct calcium regulation of TRPM7 mechanosensitive channels at plasma membrane microdomains visualized by FRET-based single cell imaging. Sci Rep 11, 17893 (2021).

30. Katanosaka, K., Takatsu, S., Mizumura, K., Naruse, K. & Katanosaka, Y. TRPV2 is required for mechanical nociception and the stretch-evoked response of primary sensory neurons. Sci Rep 8, 16782 (2018).

31. Kubanek, J., Shukla, P., Das, A., Baccus, S. A. & Goodman, M. B. Ultrasound Elicits Behavioral Responses through Mechanical Effects on Neurons and Ion Channels in a Simple Nervous System. The Journal of Neuroscience 38, 3081–3091 (2018).

32. Kubanek, J. et al. Ultrasound modulates ion channel currents. Sci Rep 6, 1–14 (2016).

33. Legon, W. et al. Transcranial focused ultrasound modulates the activity of primary somatosensory cortex in humans. Nat Neurosci 17, 322–329 (2014).

34. Murphy, K. R. et al. A tool for monitoring cell type–specific focused ultrasound neuromodulation and control of chronic epilepsy. Proc Natl Acad Sci U S A 119, e2206828119 (2022).

35. Tseng, H. et al. Region-specific effects of ultrasound on individual neurons in the awake mammalian brain. iScience 24, 102955 (2021).

36. Verhagen, L. et al. Offline impact of transcranial focused ultrasound on cortical activation in primates. Elife 8, (2019).

37. Yu, K., Niu, X., Krook-Magnuson, E. & He, B. Intrinsic functional neuron-type selectivity of transcranial focused ultrasound neuromodulation. Nat Commun 12, 2519 (2021).

38. Lowet, E. et al. Deep brain stimulation creates informational lesion through membrane depolarization in mouse hippocampus. Nat Commun 13, 7709 (2022).

39. Boraud, T., Brown, P., Goldberg, J. A., Graybiel, A. M. & Magill, P. J. Oscillations in the Basal Ganglia: The good, the bad, and the unexpected. in *The Basal Ganglia VIII* 1–24 (Kluwer Academic Publishers, Boston, 2005). doi:10.1007/0-387-28066-9_1.

40. Martinez-Losa, M. et al. Nav1.1-Overexpressing Interneuron Transplants Restore Brain Rhythms and Cognition in a Mouse Model of Alzheimer’s Disease. Neuron 98, 75–89.e5 (2018).

41. Sohal, V. S., Zhang, F., Yizhar, O. & Deisseroth, K. Parvalbumin neurons and gamma rhythms enhance cortical circuit performance. Nature 459, 698–702 (2009).

42. Park, M. et al. Effects of transcranial ultrasound stimulation pulsed at 40 Hz on Aβ plaques and brain rhythms in 5×FAD mice. Transl Neurodegener 10, 48 (2021).

43. Bobola, M. S. et al. Transcranial focused ultrasound, pulsed at 40 Hz, activates microglia acutely and reduces Aβ load chronically, as demonstrated in vivo. Brain Stimul 13, 1014–1023 (2020).

44. Iaccarino, H. F. et al. Gamma frequency entrainment attenuates amyloid load and modifies microglia. Nature 540, 230–235 (2016).

45. Lein, E. S. et al. Genome-wide atlas of gene expression in the adult mouse brain. Nature 445, 168–176 (2007).

46. Bakken, T. E. et al. Comparative cellular analysis of motor cortex in human, marmoset and mouse. Nature 598, 111–119 (2021).

47. Hodge, R. D. et al. Conserved cell types with divergent features in human versus mouse cortex. Nature 573, 61–68 (2019).

48. Tasic, B. et al. Shared and distinct transcriptomic cell types across neocortical areas. Nature 563, 72–78 (2018).

49. Dana, H. et al. High-performance calcium sensors for imaging activity in neuronal populations and microcompartments. Nat Methods 16, 649–657 (2019).

50. Kim, H. et al. Miniature ultrasound ring array transducers for transcranial ultrasound neuromodulation of freely-moving small animals. Brain Stimul 12, 251–255 (2019).

51. Ramachandran, S., Niu, X., Yu, K. & He, B. Transcranial ultrasound neuromodulation induces neuronal correlation change in the rat somatosensory cortex. J Neural Eng 19, 056002 (2022).

52. Song, K. II et al. Localization of ultrasound waveform for low intensity ultrasound-induced neuromodulation in a mouse model. *Proceedings of the Annual International Conference of the IEEE Engineering in Medicine and Biology Society*, EMBS 1122–1125 (2017) doi:10.1109/EMBC.2017.8037026.

53. Tufail, Y., Yoshihiro, A., Pati, S., Li, M. M. & Tyler, W. J. Ultrasonic neuromodulation by brain stimulation with transcranial ultrasound. Nat Protoc 6, 1453–1470 (2011).

54. Chan, E., Kovacevíc, N., Ho, S. K. Y., Henkelman, R. M. & Henderson, J. T. Development of a high resolution three-dimensional surgical atlas of the murine head for strains 129S1/SvImJ and C57Bl/6J using magnetic resonance imaging and micro-computed tomography. Neuroscience 144, 604–615 (2007).

55. Shemesh, O. A. et al. Precision Calcium Imaging of Dense Neural Populations via a Cell-Body-Targeted Calcium Indicator. Neuron 107, 470–486.e11 (2020).

56. DeFelipe, J. & Fariñas, I. The pyramidal neuron of the cerebral cortex: Morphological and chemical characteristics of the synaptic inputs. Prog Neurobiol 39, 563–607 (1992).

57. Sahara, S., Yanagawa, Y., O’Leary, D. D. M. & Stevens, C. F. The Fraction of Cortical GABAergic Neurons Is Constant from Near the Start of Cortical Neurogenesis to Adulthood. The Journal of Neuroscience 32, 4755–4761 (2012).

58. Keller, D., Erö, C. & Markram, H. Cell Densities in the Mouse Brain: A Systematic Review. Front Neuroanat 12, (2018).

59. Burks, S. R., Lorsung, R. M., Nagle, M. E., Tu, T.-W. & Frank, J. A. Focused ultrasound activates voltage-gated calcium channels through depolarizing TRPC1 sodium currents in kidney and skeletal muscle. Theranostics 9, 5517–5531 (2019).

60. Sato, T., Shapiro, M. G. & Tsao, D. Y. Ultrasonic Neuromodulation Causes Widespread Cortical Activation via an Indirect Auditory Mechanism. Neuron 98, 1031–1041.e5 (2018).

61. Guo, H. et al. Ultrasound Produces Extensive Brain Activation via a Cochlear Pathway. Neuron 98, 1020–1030.e4 (2018).

62. Mohammadjavadi, M. et al. Elimination of peripheral auditory pathway activation does not affect motor responses from ultrasound neuromodulation. Brain Stimul 12, 901–910 (2019).

63. Tyler, W. J. et al. Remote Excitation of Neuronal Circuits Using Low-Intensity, Low-Frequency Ultrasound. PLoS One 3, e3511 (2008).

64. Yang, P.-F. et al. Bidirectional and state-dependent modulation of brain activity by transcranial focused ultrasound in non-human primates. Brain Stimul 14, 261–272 (2021).

65. Gritton, H. J. et al. Unique contributions of parvalbumin and cholinergic interneurons in organizing striatal networks during movement. Nat Neurosci 22, 586–597 (2019).

66. Szabo, T. L. Causal theories and data for acoustic attenuation obeying a frequency power law. J Acoust Soc Am 97, 14–24 (1995).

67. Szabo Thomas. Diagnostic Ultrasound Imaging: Inside Out. (Elsevier, 2014). doi:10.1016/C2011-0-07261-7.

68. White, P. J., Clement, G. T. & Hynynen, K. Longitudinal and shear mode ultrasound propagation in human skull bone. Ultrasound Med Biol 32, 1085–1096 (2006).

69. George William Clarkson Kaye & Thomas Howell Laby. Tables of Physical and Chemical Constants. (Longman Publishing Group, 1995).

