## Supplemental Figures for "Ultrasound pulse repetition frequency preferentially activates different neuron populations independent of cell type"

**Supplemental Material**

**Table 1: Mechanosensitive channels in the mammalian brain.** (* indicates the ones that have been experimentally tested in contributing to ultrasound-evoked neural responses)

| **Channel Family** | **Channel Name** | **Gene Name** |
| --- | --- | --- |
| **K2P** (Two-pore potassium channel) | TREK-1* ^20,21,32^ | Kcnk2 |
|  | TREK-2*^20,32^ | Kcnk10 |
|  | TRAAK* ^20-22,32^ | Kcnk4 |
| **TRP (**transient receptor potential) | TRPC1*^19,27^, 5^27,28^, 6^27^ | Trpc 1,5,6 |
|  | TRPM2*^26^, 3^27^,7^27,29^ | Trpm 2,3,7 |
|  | TRPP1*^19^, 2*^19^ | Pkd1, 2 |
|  | TRPV2^27,30^, 4^20,27^ | Trpv 2,4 |
| **TMEM63 (**hyperosmolality-gated calcium-permeable) | TMEM63B^20^ | Tmem63B |
| **Piezo (**non-selective cationic mechanotranducer) | PIEZO1*^20,24,25^ | Piezo1 |

**Table 2: Brain-region dependent expression of mechanosensitive channels implicated in ultrasound neuromodulation. RNA In-Situ hybridization (ISH) data is from Allen Brain Institute’s Mouse Brain Atlas^48^** <https://mouse.brain-map.org>. Values are ISH [log2]. If data available from both hemispheres, they are both listed. ICTX: isocortex; OLF: olfactory areas; HPF: hippocampal formation; CTXsp: Cortical subplate; STR: striatum; PAL: Pallidum; TH: thalamus; HY: hypothalamus; MB: Midbrain; P: pons; MY: Medulla; CB: Cerebellum.

| **Channel Protein** | **Gene** | **ICTX** | **OLF** | **HPF** | **CTXsp** | **STR** | **PAL** | **TH** | **HY** | **MB** | **P** | **MY** | **CB** |
| --- | --- | --- | --- | --- | --- | --- | --- | --- | --- | --- | --- | --- | --- |
| **TREK1** | **Kcnk2** | 1.69, 1.99 | 0.94, 1.01 | 0.71, 0.80 | 1.60,  2.05 | 2.54,  2.73 | 0.57,  0.88 | 0 | 0.18,  0.33 | 0 | 0 | 0 | 0 |
| **TREK2** | **KCNK10** | 0, 1.34 | 0, 1.32 | 0, 1.48 | 0,  1.29 | 0-0.24 | 0 | 0 | 0 | 0 | 0 | 0 | 0 |
| **TRAAK** | **KCNK4** | 0 | 0 | 0 | 0 | 0 | 0 | 0 | 0 | 0 | 0 | 0 | 0 |
| **TRPC 1** | **TRPC 1** | 2.97 | 1.80 | 2.66 | 3.01 | 2.62 | 1.62 | 2.04 | 0.49 | 0.94 | 1.04 | 1.36 | 1.47 |
| **TRPC 5** | **TRPC 5** | 0, 0.43 | 0, 0.47 | 0, 1.64 | 0.76,  1.19 | 0 | 0-0.40 | 0 | 0,  0.50 | 0 | 0 | 0 | 0 |
| **TRPC 6** | **TRPC 6** | 0.60, 1.20 | 0, 0.43 | 1.92, 1.99 | 0,  0.98 | 0 | 0 | 0 | 0 | 0 | 0 | 0 | 0 |
| **TRPM 2** | **TRPM 2** | 1.75 | 1.56 | 1.41 | 1.89 | 1.06 | 1.67 | 2.44 | 1.38 | 1.99 | 1.71 | 2.21 | 2.21 |
| **TRPM 3** | **TRPM 3** | 0 | 1.05 | 1.70 | 0.38 | 0 | 0 | 0 | 0 | 0 | 0 | 0 | 0 |
| **TRPM 7** | **TRPM 7** | 1.96 | 1.80 | 1.68 | 1.50 | 1.03 | 0.89 | 1.02 | 1.01 | 1.31 | 0.87 | 1.06 | 0.44 |
| **TRPV 2** | **TRPV 2** | 0 | 0 | 0 | 0 | 0 | 0 | 0 | 0 | 0 | 0 | 0 | 0 |
| **TRPV 4** | **TRPV 4** | 0 | 0 | 0 | 0 | 0 | 0 | 0 | 0 | 0 | 0 | 0 | 0 |
| **TMEM63B** | **TMEM63B** | 2.89 | 2.10 | 2.57 | 2.38 | 1.40 | 1.87 | 2.41 | 1.76 | 2.11 | 1.59 | 1.96 | 1.22 |
| **Piezo1** | **Piezo1** | 3.58 | 3.31 | 3.76 | 3.66 | 2.92 | 2.8 | 3.14 | 3.22 | 3.21 | 2.68 | 2.87 | 4.39 |

**Table 3: Differential gene expression in glutamatergic excitatory neurons versus parvalbumin-positive interneurons in the human motor cortex.** (Data is from <https://viewer.cytosplore.org/>)

| **Channel Protein** | **Gene** | **Mean expression glutamatergic** | **Mean expression**  **Pvalb** | **Differential expression (glutamatergic-Pvalb)** | **Bonferroni adjusted p value** |
| --- | --- | --- | --- | --- | --- |
| **KCNK10** | **Kcnk10** | 3.759 | 2.121 | 1.638 | 3.58E-09 |
| **TRPM3** | **Trpm3** | 7.183 | 5.845 | 1.338 | 1.53E-99 |
| **KCNK4** | **Kcnk4** | 1.278 | 0.256 | 1.022 | 1 |
| **TRPC6** | **Trpc6** | 1.407 | 0.410 | 0.997 | 4.48E-03 |
| **TRPM7** | **Trpm7** | 6.599 | 6.032 | 0.567 | 4.56E-59 |
| **TRPV2** | **Trpv2** | 0.485 | 0.152 | 0.333 | 1 |
| **PIEZO1** | **Piezo1** | 0.069 | 0.037 | 0.032 | 1 |
| **TRPV4** | **Trpv4** | 0.057 | 0.059 | -0.002 | 1 |
| **TMEM63B** | **Tmem63b** | 2.929 | 3.216 | -0.287 | 1.04E-98 |
| **KCNK2** | **Kcnk2** | 1.512 | 1.933 | -0.421 | 3.16E-32 |
| **TRPM2** | **Trpm2** | 4.446 | 4.954 | -0.508 | 1.83E-119 |
| **TRPC1** | **Trpc1** | 6.926 | 7.701 | -0.776 | 0 |
| **TRPC5** | **Trpc5** | 2.962 | 6.646 | -3.685 | 0 |

**Table 4: Differential gene expression in glutamatergic excitatory neurons and GABAergic interneurons in the mouse motor cortex and hippocampus^46-48^.** (Data is from <https://viewer.cytosplore.org/>)

| **Channel Protein** | **Gene** | **Mean expression**  **glutamatergic** | **Mean expression**  **GABAergic** | **Mean differential**  **expression** | **Bonferroni**  **adjusted p value** |
| --- | --- | --- | --- | --- | --- |
| **TRAAK** | **Kcnk4** | 3.528 | 0.835 | 2.693 | 0 |
| **TRPM 3** | **Trpm3** | 3.388 | 1.772 | 1.616 | 2.30E-65 |
| **TREK2** | **Kcnk10** | 2.123 | 0.593 | 1.530 | 9.58E-44 |
| **TRPC 1** | **Trpc1** | 4.000 | 2.865 | 1.134 | 5.60E-58 |
| **TREK1** | **Kcnk2** | 3.920 | 2.963 | 0.957 | 1.89E-294 |
| **TRPV 2** | **Trpv2** | 2.815 | 2.164 | 0.651 | 3.02E-12 |
| **TRPM 7** | **Trpm7** | 4.977 | 4.484 | 0.494 | 1 |
| **TMEM63B** | **Tmem63b** | 4.540 | 4.048 | 0.492 | 1 |
| **TRPC 5** | **Trpc5** | 2.682 | 2.510 | 0.171 | 4.80E-147 |
| **TRPV 4** | **Trpv4** | 0.024 | 0.026 | -0.002 | 1 |
| **Piezo1** | **Piezo1** | 0.183 | 0.238 | -0.055 | 1.90E-27 |
| **TRPM 2** | **Trpm2** | 1.767 | 1.871 | -0.104 | 9.80E-30 |
| **TRPC 6** | **Trpc6** | 1.634 | 2.216 | -0.582 | 0 |

**Table 5: Experimental pulse timing parameters**

| **Stimulation regime** | **Type** | **Duration** | **Ramp duration** | **Ramp shape** | **Repetition interval / frequency** |
| --- | --- | --- | --- | --- | --- |
| 10 Hz,  single PRF | Pulse | 20 ms | 0 s | rectangular | 100 ms/10 Hz |
| 10 Hz,  single PRF | Pulse train | 1 s | 0 s | rectangular |  |
| 40 Hz,  single PRF | Pulse | 8 ms | 0 s | rectangular | 25 ms/40 Hz |
| 40 Hz,  single PRF | Pulse train | 1 s | 0 s | rectangular |  |
| 140 Hz,  single PRF | Pulse | 1.43 ms | 0 s | rectangular | 7.14 ms/140 Hz |
| 140 Hz,  single PRF | Pulse train | 1 s | 0 s | rectangular |  |
| 2000 Hz,  single PRF | Pulse | 0.22 ms | 0 s | rectangular | 0.5 ms/2 kHz |
| 2000 Hz,  single PRF | Pulse train | 1 s | 0 s | rectangular |  |
| 10 Hz,  alternating PRF | Pulse | 20 ms | 0 s | rectangular | 100 ms/10 Hz |
| 10 Hz,  alternating PRF | Pulse train | 1 s | 0 s | rectangular |  |
| 40 Hz,  alternating PRF | Pulse | 8 ms | 0 s | rectangular | 25 ms/40 Hz |
| 40 Hz,  alternating PRF | Pulse train | 1 s | 0 s | rectangular |  |
| 140 Hz, alternating PRF | Pulse | 1.43 ms | 0 s | rectangular | 7.14 ms/140 Hz |
| 140 Hz, alternating PRF | Pulse train | 1 s | 0 s | rectangular |  |

**Table 6: Acoustic safety parameters**

| **Frequency**  **/PRF** | **Spatial peak pressure** | **I_SPPA_ (W/cm^2^)** | **I_SPTA_ (W/cm^2^)** | **Total acoustic cycles** | **Thermal Index (TIC)** | ***In situ***  **spatial peak pressure** | ***In situ* peak neg. pressure** | **Mechanical Index** |
| --- | --- | --- | --- | --- | --- | --- | --- | --- |
| 350 kHz/  10 Hz | 522 kPa | 9.11 | 1.82 | 70,000 | 0.18 | 289 kPa | -289 kPa | 0.49 |
| 350 kHz/  40 Hz | 522 kPa | 9.11 | 1.82 | 70,000 | 0.18 | 289 kPa | -289 kPa | 0.49 |
| 350 kHz/  140 Hz | 522 kPa | 9.11 | 1.82 | 70,000 | 0.18 | 289 kPa | -289 kPa | 0.49 |
| 350 kHz/  2 kHz | 522 kPa | 9.11 | 3.91 | 150,000 | 0.39 | 289 kPa | -289 kPa | 0.49 |

**Table 7: Ultrasound stimulation efficacy**

| **PRF/**  **Mouse ID** | **Total Neurons**  **(#)** | **Modulated Neurons**  **(#)** | **Percent Modulated**  **(%)** |
| --- | --- | --- | --- |
| **10 Hz (20% DC)** | **861** | **162** | **18.82%** |
| 611276 | 120 | 42 | 35.00% |
| 611282 | 284 | 10 | 3.52% |
| 611302 | 353 | 98 | 27.76% |
| 617456 | 104 | 12 | 11.54% |
| **40 Hz (20% DC)** | **281** | **22** | **7.83%** |
| 31554 | 177 | 17 | 9.60% |
| 611282 | 104 | 5 | 4.81% |
| **140 Hz (20% DC)** | **1019** | **183** | **17.96%** |
| 611282 | 292 | 91 | 31.16% |
| 611302 | 416 | 60 | 14.42% |
| 617456 | 311 | 32 | 10.29% |
| **2000 Hz (42% DC)** | **1192** | **200** | **16.78%** |
| 615295 | 230 | 65 | 28.26% |
| 617435 | 488 | 75 | 15.37% |
| 617456 | 474 | 60 | 12.66% |
| **Grand Total** | **3353** | **567** | **16.91%** |

**Table 8: Modulation by sham stimulation**

| **Sham PRF/**  **Mouse ID** | **Total Neurons**  **(#)** | **Modulated Neurons**  **(#)** | **Percent Modulated**  **(%)** |
| --- | --- | --- | --- |
| **10 Hz** | **431** | **17** | **3.94%** |
| 31557 | 217 | 10 | 4.61% |
| 31624 | 61 | 3 | 4.92% |
| 611276 | 59 | 2 | 3.39% |
| 611277 | 49 | 1 | 2.04% |
| 611282 | 45 | 1 | 2.22% |
| **40 Hz** | **458** | **36** | **7.86%** |
| 31557 | 233 | 21 | 9.01% |
| 31624 | 95 | 7 | 7.37% |
| 611276 | 64 | 6 | 9.38% |
| 611277 | 24 | 1 | 4.17% |
| 611282 | 42 | 1 | 2.38% |
| **140** | **726** | **20** | **2.75%** |
| 31554 | 97 | 7 | 7.22% |
| 31557 | 213 | 2 | 0.94% |
| 31624 | 281 | 5 | 1.78% |
| 611276 | 57 | 5 | 8.77% |
| 611277 | 28 | 0 | 0.00% |
| 611282 | 50 | 1 | 2.00% |
| **2000** | **300** | **14** | **4.67%** |
| 31544 | 130 | 7 | 5.38% |
| 31557 | 92 | 6 | 6.52% |
| 50354 | 78 | 1 | 1.28% |
| **Grand Total** | **1915** | **87** | **4.54%** |

**Table 9: PV cell stimulation overview**

| **Mouse /**  **Session** | **Total (#)** | **PV**  **(#)** | **10 Hz (#)** | **10 Hz (%)** | **40 Hz (#)** | **40 Hz (%)** | **140 Hz (#)** | **140 Hz (%)** |
| --- | --- | --- | --- | --- | --- | --- | --- | --- |
| **78596** | **709** | **125** | **12** | **9.60%** | **15** | **12.00%** | **6** | **4.80%** |
| 1 | 213 | 34 | 2 | 5.88% | 2 | 5.88% | 1 | 2.94% |
| 2 | 240 | 37 | 3 | 8.11% | 7 | 18.92% | 2 | 5.41% |
| 3 | 115 | 26 | 4 | 15.38% | 2 | 7.69% | 1 | 3.85% |
| 4 | 141 | 28 | 3 | 10.71% | 4 | 14.29% | 2 | 7.14% |
| **81623** | **296** | **29** | **5** | **17.24%** | **6** | **20.69%** | **10** | **34.48%** |
| 1 | 106 | 10 | 3 | 30.00% | 1 | 10.00% | 2 | 20.00% |
| 2 | 96 | 7 | 0 | 0.00% | 1 | 14.29% | 3 | 42.86% |
| 3 | 30 | 2 | 0 | 0.00% | 0 | 0.00% | 0 | 0.00% |
| 4 | 64 | 10 | 2 | 20.00% | 4 | 40.00% | 5 | 50.00% |
| **81625** | **347** | **40** | **3** | **7.50%** | **7** | **17.50%** | **5** | **12.50%** |
| 1 | 71 | 5 | 1 | 20.00% | 1 | 20.00% | 0 | 0.00% |
| 2 | 77 | 7 | 0 | 0.00% | 2 | 28.57% | 0 | 0.00% |
| 3 | 72 | 12 | 0 | 0.00% | 1 | 8.33% | 2 | 16.67% |
| 4 | 127 | 16 | 2 | 12.50% | 3 | 18.75% | 3 | 18.75% |
| **81627** | **249** | **26** | **2** | **7.69%** | **4** | **15.38%** | **2** | **7.69%** |
| 1 | 18 | 0 | 0 | -- | 0 | -- | 0 | -- |
| 2 | 123 | 16 | 2 | 12.50% | 2 | 12.50% | 1 | 6.25% |
| 3 | 108 | 10 | 0 | 0.00% | 2 | 20.00% | 1 | 10.00% |
| **81645** | **271** | **19** | **6** | **31.58%** | **5** | **26.32%** | **3** | **15.79%** |
| 1 | 40 | 2 | 1 | 50.00% | 0 | 0.00% | 1 | 50.00% |
| 2 | 103 | 11 | 4 | 36.36% | 3 | 27.27% | 2 | 18.18% |
| 3 | 128 | 6 | 1 | 16.67% | 2 | 33.33% | 0 | 0.00% |
| **81646** | **340** | **30** | **8** | **26.67%** | **6** | **20.00%** | **4** | **13.33%** |
| 1 | 104 | 16 | 3 | 18.75% | 2 | 12.50% | 3 | 18.75% |
| 2 | 112 | 8 | 3 | 37.50% | 2 | 25.00% | 1 | 12.50% |
| 3 | 124 | 6 | 2 | 33.33% | 2 | 33.33% | 0 | 0.00% |
| **Grand Total** | **2212** | **269** | **36** | **13.38%** | **43** | **15.99%** | **30** | **11.15%** |

**Table 10: Non-PV cell stimulation overview**

| **Mouse/**  **Session** | **Total (#)** | **Non-PV**  **(#)** | **10 Hz (#)** | **10 Hz (%)** | **40 Hz (#)** | **40 Hz (%)** | **140 Hz (#)** | **140 Hz (%)** |
| --- | --- | --- | --- | --- | --- | --- | --- | --- |
| **78596** | **709** | **584** | **61** | **10.45%** | **43** | **7.36%** | **60** | **10.27%** |
| 1 | 213 | 179 | 4 | 2.23% | 9 | 5.03% | 13 | 7.26% |
| 2 | 240 | 203 | 26 | 12.81% | 16 | 7.88% | 17 | 8.37% |
| 3 | 115 | 89 | 5 | 5.62% | 2 | 2.25% | 13 | 14.61% |
| 4 | 141 | 113 | 26 | 23.01% | 16 | 14.16% | 17 | 15.04% |
| **81623** | **296** | **267** | **48** | **17.98%** | **46** | **17.23%** | **42** | **15.73%** |
| 1 | 106 | 96 | 19 | 19.79% | 24 | 25.00% | 8 | 8.33% |
| 2 | 96 | 89 | 11 | 12.36% | 11 | 12.36% | 11 | 12.36% |
| 3 | 30 | 28 | 5 | 17.86% | 1 | 3.57% | 2 | 7.14% |
| 4 | 64 | 54 | 13 | 24.07% | 10 | 18.52% | 21 | 38.89% |
| **81625** | **347** | **307** | **40** | **13.03%** | **50** | **16.29%** | **37** | **12.05%** |
| 1 | 71 | 66 | 13 | 19.70% | 13 | 19.70% | 8 | 12.12% |
| 2 | 77 | 70 | 15 | 21.43% | 19 | 27.14% | 10 | 14.29% |
| 3 | 72 | 60 | 5 | 8.33% | 8 | 13.33% | 7 | 11.67% |
| 4 | 127 | 111 | 7 | 6.31% | 10 | 9.01% | 12 | 10.81% |
| **81627** | **249** | **223** | **21** | **9.42%** | **28** | **12.56%** | **24** | **10.76%** |
| 1 | 18 | 18 | 1 | 5.56% | 7 | 38.89% | 3 | 16.67% |
| 2 | 123 | 107 | 11 | 10.28% | 7 | 6.54% | 12 | 11.21% |
| 3 | 108 | 98 | 9 | 9.18% | 14 | 14.29% | 9 | 9.18% |
| **81645** | **271** | **252** | **47** | **18.65%** | **36** | **14.29%** | **31** | **12.30%** |
| 1 | 40 | 38 | 5 | 13.16% | 5 | 13.16% | 3 | 7.89% |
| 2 | 103 | 92 | 26 | 28.26% | 22 | 23.91% | 16 | 17.39% |
| 3 | 128 | 122 | 16 | 13.11% | 9 | 7.38% | 12 | 9.84% |
| **81646** | **340** | **310** | **54** | **17.42%** | **57** | **18.39%** | **31** | **10.00%** |
| 1 | 104 | 88 | 13 | 14.77% | 13 | 14.77% | 10 | 11.36% |
| 2 | 112 | 104 | 24 | 23.08% | 19 | 18.27% | 10 | 9.62% |
| 3 | 124 | 118 | 17 | 14.41% | 25 | 21.19% | 11 | 9.32% |
| **Grand Total** | **2212** | **1943** | **271** | **13.95%** | **260** | **13.38%** | **225** | **11.58%** |

**Table 11: Sham stimulation overview**

| **Mouse/**  **Session** | **Total**  **(#)** | **10 Hz**  **(#)** | **10 Hz**  **(%)** | **40 Hz**  **(#)** | **40 Hz**  **(%)** | **140 Hz**  **(#)** | **140 Hz**  **(%)** |
| --- | --- | --- | --- | --- | --- | --- | --- |
| **78596** | **761** | **39** | **5.12%** | **22** | **2.89%** | **15** | **1.97%** |
| 1 | 302 | 5 | 1.66% | 9 | 2.98% | 4 | 1.32% |
| 2 | 238 | 20 | 8.40% | 8 | 3.36% | 6 | 2.52% |
| 3 | 221 | 14 | 6.33% | 5 | 2.26% | 5 | 2.26% |
| **81623** | **258** | **9** | **3.49%** | **10** | **3.88%** | **8** | **3.10%** |
| 1 | 184 | 5 | 2.72% | 6 | 3.26% | 6 | 3.26% |
| 2 | 74 | 4 | 5.41% | 4 | 5.41% | 2 | 2.70% |
| **81625** | **334** | **12** | **3.59%** | **15** | **4.49%** | **15** | **4.49%** |
| 1 | 108 | 4 | 3.70% | 4 | 3.70% | 4 | 3.70% |
| 2 | 94 | 2 | 2.13% | 4 | 4.26% | 4 | 4.26% |
| 3 | 132 | 6 | 4.55% | 7 | 5.30% | 7 | 5.30% |
| **81627** | **90** | **7** | **7.78%** | **3** | **3.33%** | **6** | **6.67%** |
| 1 | 90 | 7 | 7.78% | 3 | 3.33% | 6 | 6.67% |
| **81645** | **268** | **10** | **3.73%** | **14** | **5.22%** | **12** | **4.48%** |
| 1 | 152 | 8 | 5.26% | 3 | 1.97% | 9 | 5.92% |
| 2 | 116 | 2 | 1.72% | 11 | 9.48% | 3 | 2.59% |
| **81646** | **148** | **8** | **5.41%** | **3** | **2.03%** | **5** | **3.38%** |
| 1 | 90 | 4 | 4.44% | 2 | 2.22% | 3 | 3.33% |
| 2 | 58 | 4 | 6.90% | 1 | 1.72% | 2 | 3.45% |
| **Grand Total** | **1859** | **85** | **4.57%** | **67** | **3.60%** | **61** | **3.28%** |

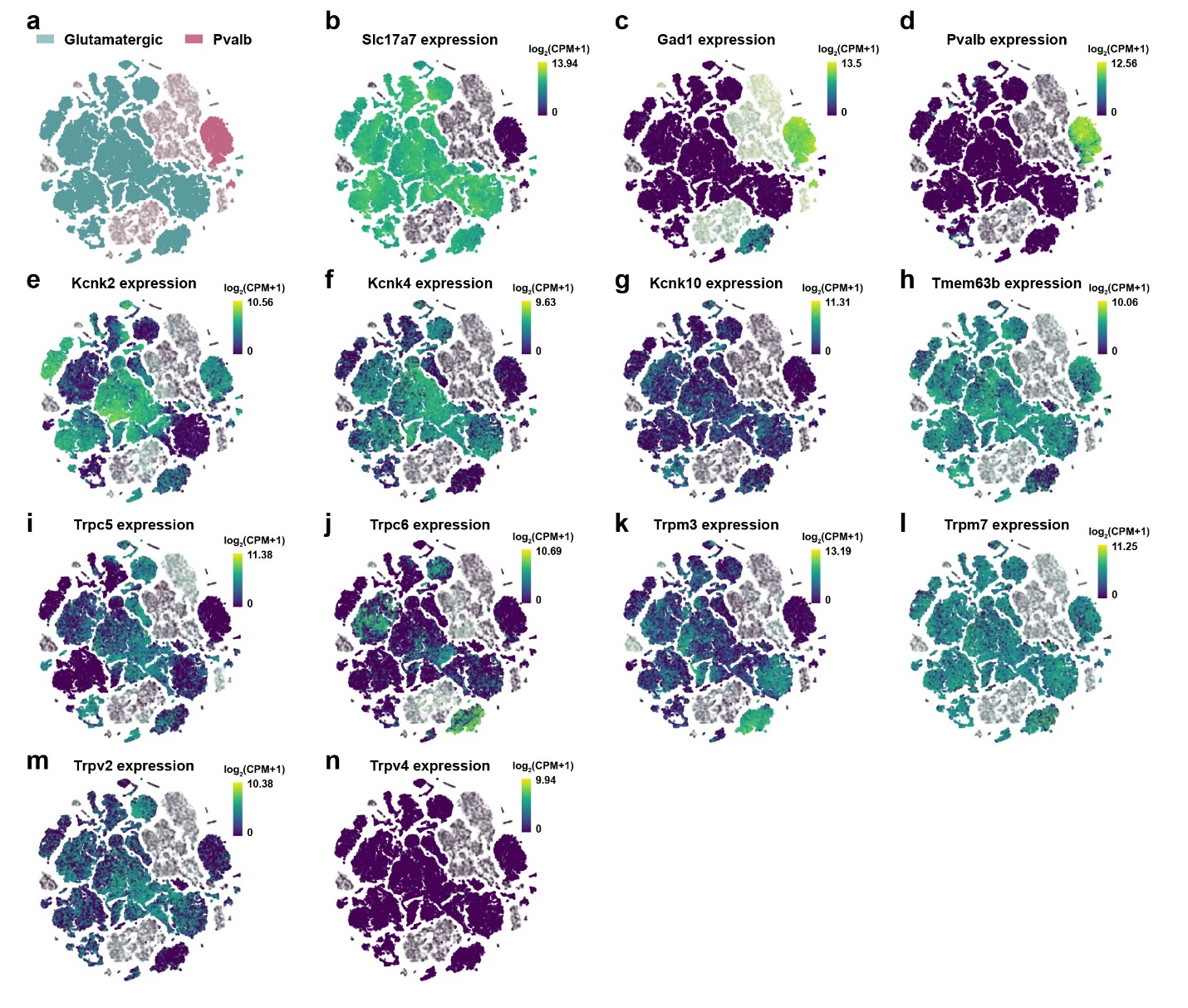

**Supplemental Figure 1: Expression levels of major mechanosensitive channels in glutamatergic excitatory neurons versus parvalbumin (PV)-positive interneurons in mouse cortex and hippocampus.** (**a-n**), Uniform Manifold Approximation and Projection (UMAP) representation of the full single-cell sequencing results, with proximal clusters representing individual cell subtypes. Single cell RNA-Seq data is from the Allen Brain Institute’s Cell Types Database (https://brain-map.org) (**a**). Glutamatergic excitatory neurons and parvalbumin-expressing (PV) interneurons are highlighted for differential expression analysis. Other cellular subtypes are blurred out for ease of comparison. (**b**). Excitatory neuron marker gene Slc17a7 expression UMAP. (**c**). Inhibitory neuron marker gene Gad1 expression UMAP. (**d**). PV neuron marker Pvalb expression UMAP. (**e-n**), UMAP for selected mechanosensitive channel genes, scaled by log_2_(CPM+1).

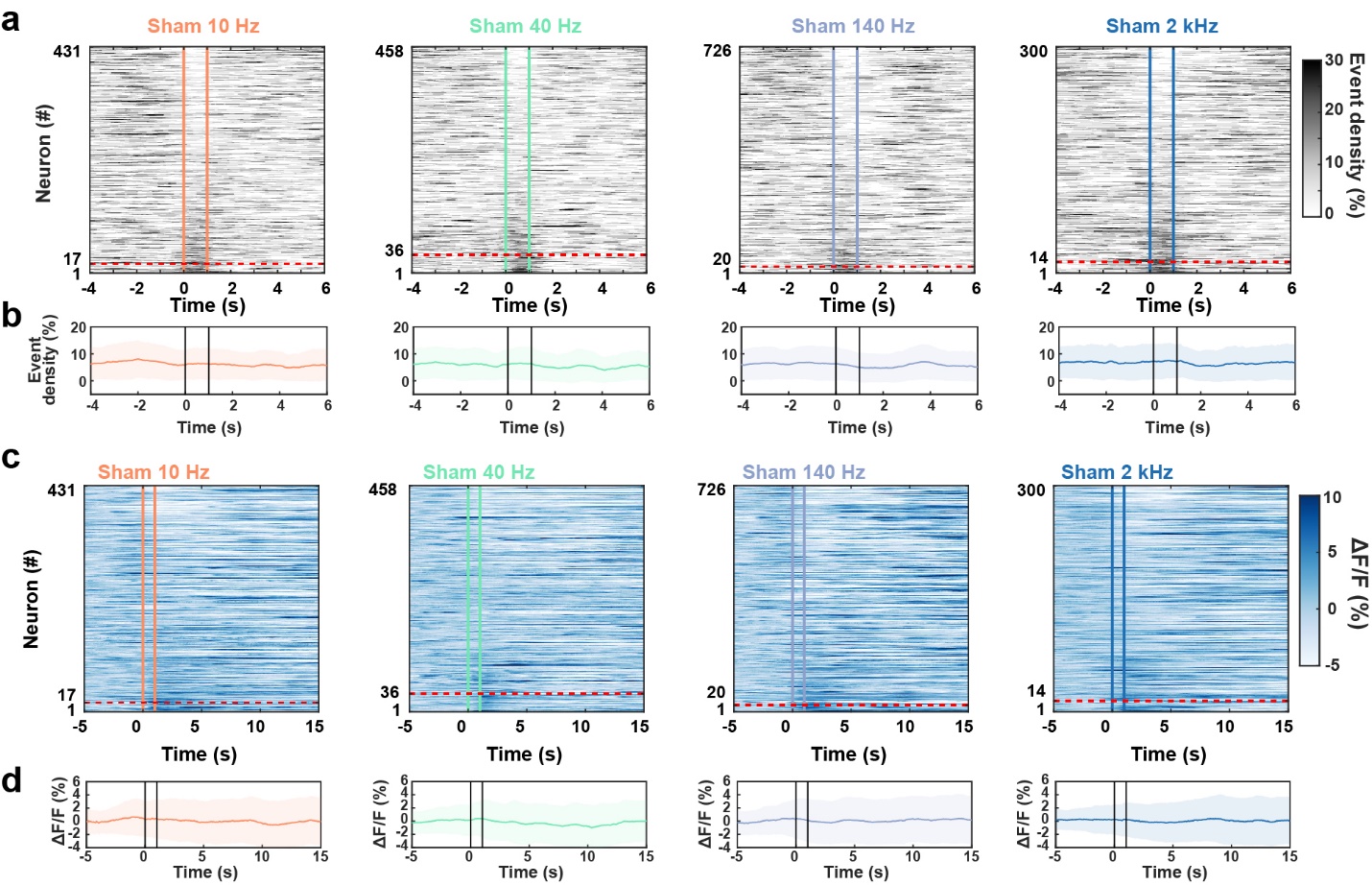

**Supplemental Figure 2:** (**a**). Population average event density changes across all neurons before, during, and after sham stimulation pulsed at various PRFs. Vertical colored lines represent sham onset and offset. Neurons were sorted by event density difference during 1 second after versus before sham onset. Neurons classified as activated by sham are below the red dashed line. (**b**). Sham population event density profiles. Shaded lines correspond to mean event density ± SD. (**c**). Normalized ΔF/F heatmaps representing average calcium fluorescence during sham stimulation. Colored lines represent sham stimulation onset and offset. Neurons were sorted by event density as in **Fig. 2a.** Sham neurons classified as activated are below the red dashed line. (**d**). Sham population ΔF/F profiles. Shaded lines correspond to mean ΔF/F ± SD.

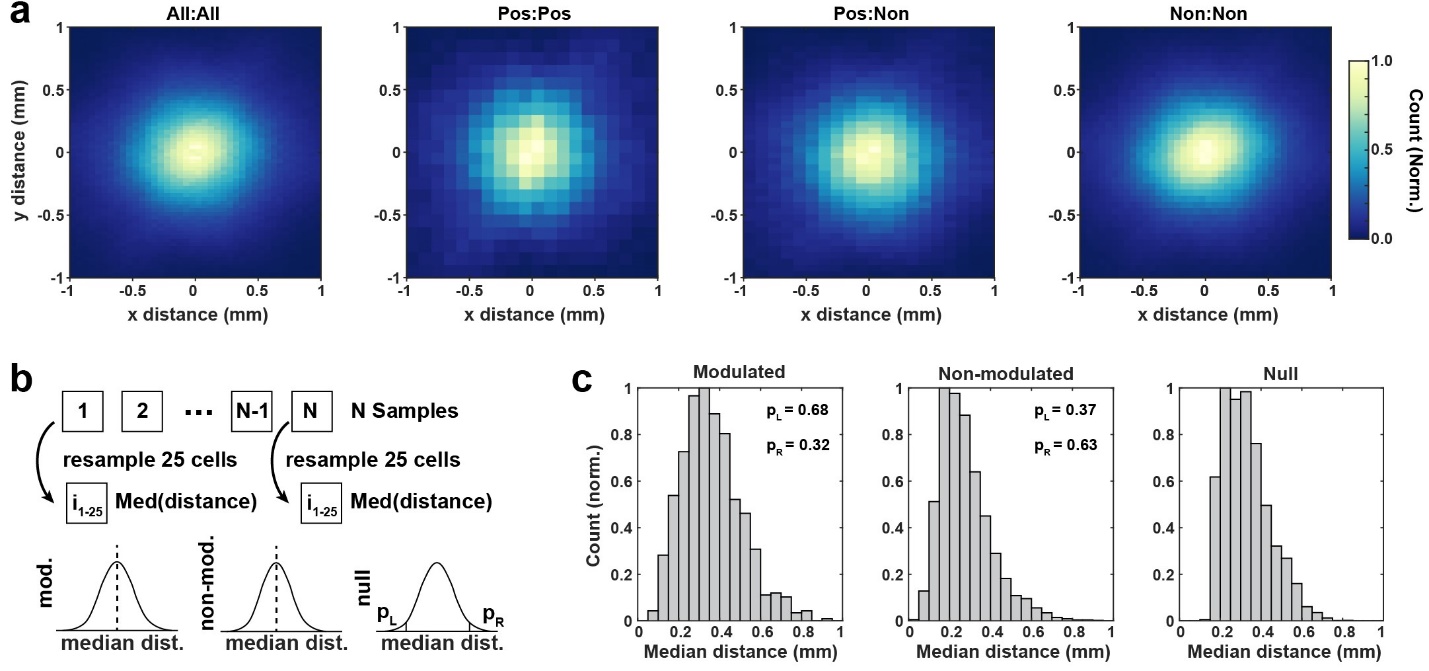

**Supplemental Figure 3: (a)**. 2D histograms of relative distance between neuron pairs across all imaging sessions. Color represents normalized count. **(b)**. Schematic of statistical resampling test for median distance of modulated and non-modulated cell populations vs the full cell population (“null”). For 1000 iterations, 25 cells were selected without replacement, and the median distance between each cell pair formed the null distribution. If any sampled cells were modulated, the median distance between those cells was calculated to form the modulated test distribution. Similarly, the non-modulated test distribution was formed from the median distance between sampled pairs of non-modulated cells. The left and right p-values for each test distribution were calculated based on the null distribution. **(c)** Neither the modulated (left) nor the non-modulated (middle) cell populations had a significantly different median distance between cells than the null distribution (right), indicating no spatial patterning of modulation (p_L,R_ > 0.05 for both groups).

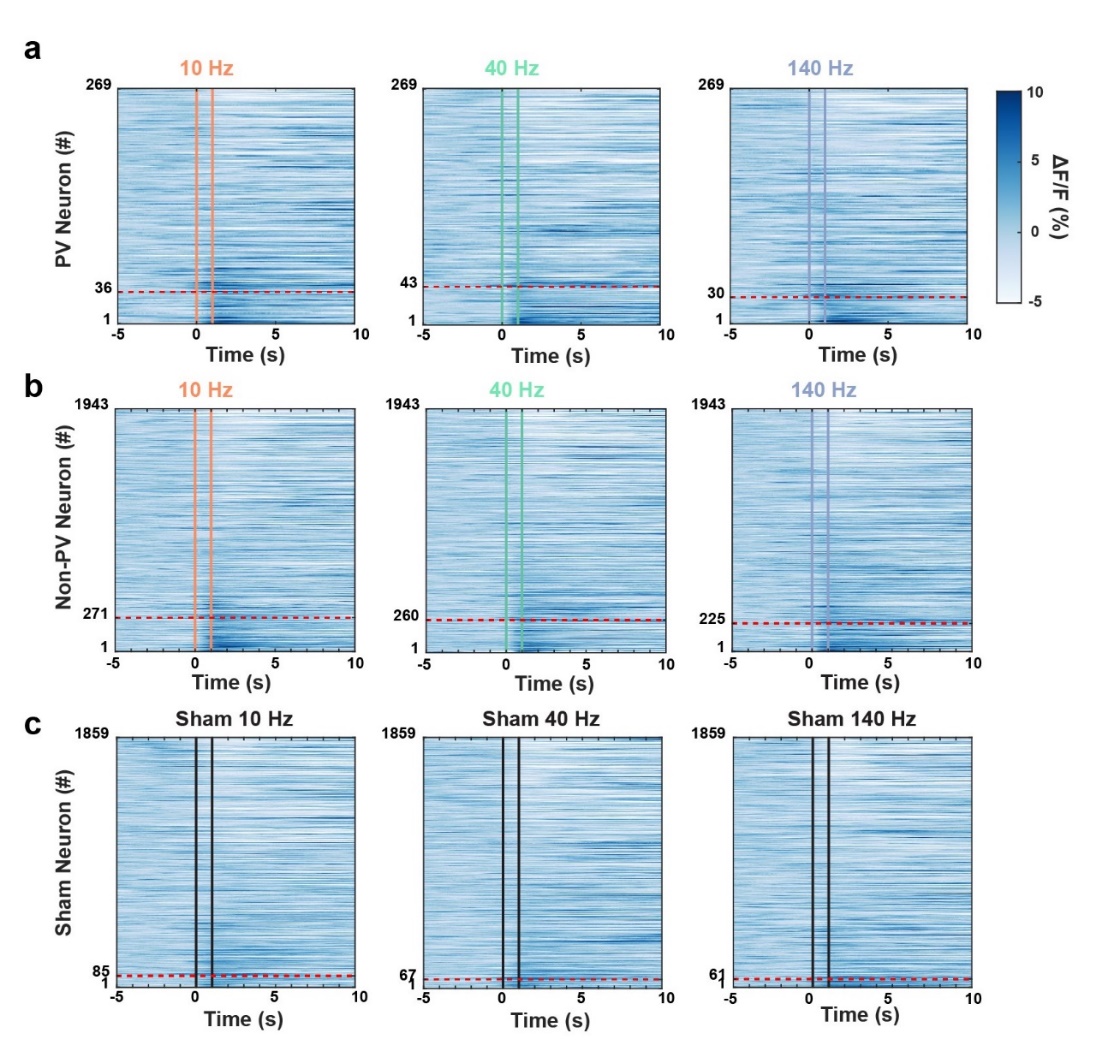

**Supplemental Figure 4:** (**a-c**) Normalized ΔF/F heatmaps across **(a)** PV neurons, **(b)** non-PV neurons, and **(c)** neurons during sham stimulation. The colored lines represent US onset and offset. Neurons were sorted by event density difference during as in **7a**. Neurons classified as activated by US or sham are below the red dashed line.

**Supplemental Figure 6**

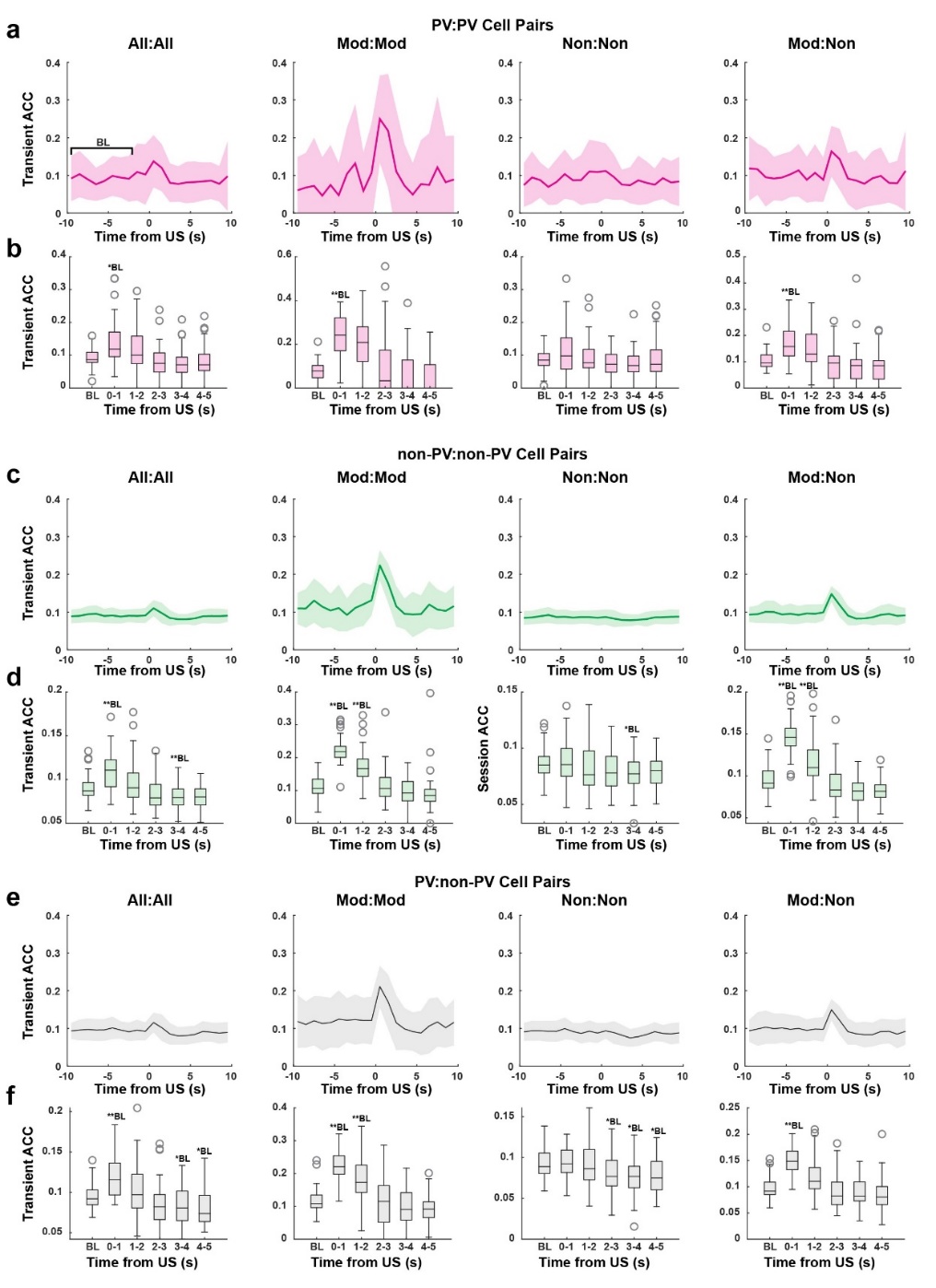

**Supplemental Figure 5:** (**a,c,e**) Time-course of ACC changes (“Transient ACC”) over each second for (**a**) PV cell pairs, (**c**) non-PV cell pairs, and (**e**) PV:non-PV cell pairs. Traces are plotted as session-wise mean ACC ± SD. Inset: bar plot with mean ACC ± SD used for statistical comparison. The first eight seconds of baseline were averaged for statistical comparison. (**b,d,f**) Statistical comparison of transient ACC values during baseline (“BL”) and post-US for (**b**) PV cell pairs, (**d**) non-PV cell pairs, and (**f**) PV:non-PV cell pairs. Friedman test was significant for all comparisons (p < 0.05). *BL and **BL represent significant Nemenyi rank difference from baseline at a given timepoint for significance levels α = 0.05 and α = 0.01, respectively.
